# Transcriptomic reprogramming in a susceptible *Phaseolus vulgaris* L. variety during *Pseudomonas syringae* attack: The key role of homogalacturonan methylation

**DOI:** 10.1101/2022.12.19.521100

**Authors:** Alfonso G. De la Rubia, Asier Largo-Gosens, Ricardo Yusta, Pablo Sepúlveda, Aníbal Riveros, Mª Luz Centeno, Dayan Sanhueza, Claudio Meneses, Susana Saez-Aguayo, Penélope García-Angulo

**Author notes:** Both authors contribute equally to this work (Alfonso G. De la Rubia, Asier Largo-Gosens). Corresponding authors: Susana Saez-Aguayo and Penélope García-Angulo. **Material distribution footnote.** “The author(s) responsible for the distribution of materials integral to the findings presented in this article in accordance with the policy described in the Instructions for Authors (https://academic.oup.com/plcell/pages/General-Instructions) are: Susana Saez-Aguayo and Penélope García-Angulo. The datasets generated and analyzed for this study can be found in the National Center for Biotechnology Information (NCBI) repository, PRJNA911667 http://www.ncbi.nlm.nih.gov/bioproject/911667.

## Abstract

The susceptibility of common bean varieties to *Pseudomonas syringae* pv. *phaseolicola* (Pph) has been well-documented. However, the molecular mechanism that drives this susceptibility has not been clarified yet. In an attempt to understand this process, 15-day-old common bean plants, variety *riñón,* were infected with Pph to analyze the transcriptomic changes during the first steps of the infection (at 2 and 9 h). RNA-seq analysis showed an upregulation of defense-and signaling-related genes at 2h, most of them being downregulated at 9h, suggesting that Pph would inhibit the transcriptomic reprogramming of the plant. This trend was also observed in the modulation of 101 cell wall (CW) related genes, suggesting that Pph could produce/induce changes in the CW. However, the changes in CW composition at early stages of Pph infection were related to homogalacturonan (HG) methylation and the formation of HG egg boxes. From all HG-related genes modulated by the infection, a common bean pectin methylesterase inhibitor 3 (*PvPMEI3*) gene – closely related to *AtPMEI3* — was detected. In addition, PMEI3 protein was located in the apoplast and its PME inhibitory activity was demonstrated. Therefore, PvPMEI3 seems to be a good candidate to play a key role in Pph infection. This premise was supported by the analysis of Arabidopsis *pmei3* mutant, which showed susceptibility to Pph, in contrast to resistant Col-0 control plants. All these changes could be an attempt to reinforce the CW structure and thus, hinder the attack of the bacterium. However, these transcriptional and CW-remodeling processes are neither maintained during the necessary time, nor are deep enough to block the action of the pathogen, facilitating the well-known susceptibility of *riñón* variety to Pph.

## Introduction

In nature, plants, as sessile organisms, are usually under biotic stress caused by different pathogens, affecting their development and, in some cases, their survival (Dangl et al., 2013). To combat attackers, plants have developed multiple defensive strategies, tightly regulated, that constitute a sophisticated immune system (Jones and Dangl, 2006; Monaghan and Zipfel, 2012; Zipfel et al., 2014).

This response begins when plants can perceive certain structures of a pathogen outside the cell by means of receptors located in the cell membrane to initiate a first defense line known as pattern-triggered immunity (PTI) (Monaghan and Zipfel, 2012). However, some pathogens can break the first line of defense of the plant, the cell wall (CW), introducing effectors into the cytoplasm to limit PTI response (Jones and Dangl, 2006). In this case, plant defense continues inside the cell by receptors that perceive pathogen-derived effectors, activating a second defensive line, named effector-triggered immunity (ETI) (Dodds and Rathjen, 2010). Both PTI and ETI lead to a transcriptional reprogramming and synthesis of complex metabolites that induct hypersensitive response (HR), a process which can be also blocked by pathogen effectors (Jones and Dangl, 2006).

In all of these defensive processes, several factors such as receptors, reactive oxygen species (ROS) production, calcium influx, phytohormones (PHs) and different signaling cascades, play important roles. Once the pathogen is perceived by the membrane receptors, some signaling pathways are activated (Couto and Zipfel, 2016;), such as ROS burst and rapid calcium ion-influx, which usually activate MAPK cascades that finally cause a transcriptional reprogramming (Couto and Zipfel, 2016; Zhou and Zhang, 2020). As in other physiological processes, PHs play an important role during the immune defense activation and its maintenance, although the events following pathogen recognition remain largely unknown (Couto and Zipfel, 2016). Among others, Salicylic acid (SA), Jasmonic acid (JA), Abscisic acid (ABA) and ethylene have been proved to be involved in different immune responses against different pathogens (Bari and Jones, 2009). The other crucial element in plant-pathogen interaction is the CW. Firstly thought as a physical barrier that hinders the entry of the pathogen into the protoplast, the CW is also the first system that participates in the perception of the intruder upon pathogen apparition (Bacete et al., 2018; Vaahtera et al., 2019; Molina et al., 2021) and it is a metabolically-active cellular compartment in which different enzymes participate reinforcing and/or maintaining the structure (Delaunois et al., 2014; Farvardin et al., 2020). This defensive structure is composed of cellulose, hemicellulose (such as xyloglucan or xylans) and pectin (such as homogalacturonan -HG- and rhamnogalacturonan I and II –RGI and RGII-) polysaccharides, whose proportions can vary depending on the tissue and plant species and under the pressure of different biotic and abiotic stresses (Baez et al., 2022). The development of disease could be the result of the pathogen interfering with one or more of all these defensive responses.

In the case of common bean variety *riñón*, which is susceptible to the attack of the biotrophic gamma-proteobacteria *Pseudomonas syringae* pv. *phaseolicola* (Pph), some of the defensive pathways, as well as some probable points where the pathogen interferes with these pathways, have already been described in De la Rubia et al. (2021a and b). It was determined that, shortly after the infection starts, this variety is able to perceive the presence of Pph but it is unable to produce a quick and effective defense response due to the lack of a peak in the production of SA soon after infection (De la Rubia et al., 2021a). In addition, the inoculation of Pph modified the pectic components of common bean CWs after 15 days of infection (De la Rubia et al., 2021b). However, when common bean immunity was treated with a SA structural analogue known as 2,6-dichloroisonicotinic acid (INA), a drastic CW remodeling was triggered, by increasing the cellulose content, hindering the extraction of the matrix polysaccharides from the CW, and increasing the molecular weight of these polysaccharides. The INA treatment increased the HG and RG cross-linking, which finally resulted in a CW more resistant to enzymatic hydrolysis. The inoculation with Pph after INA priming did not modify this CW remodeling substantially, and the resultant CWs were as resistant to enzymatic degradation as those extracted from INA-treated plants without Pph inoculation (De la Rubia et al., 2021b). Considering all these results, the modulation of pectin structure seems to be determinant in the common bean-Pph pathosystem.

The role of pectins in defense has also been attested in different works carried out by the characterization of several Arabidopsis mutants, in which a specific CW alteration induces resistance or vulnerability against different diseases (Lionetti et al., 2017; Silva-Sanzana et al., 2019; Molina et al., 2021). For example, Arabidopsis mutants *gae1* and *gae6*, repressed in glucuronate 4-epimerase genes, presented alterations in the content of galacturonic acid, producing less resistance to *Pseudomonas syringae* pv. *maculicola* and *Botrytis cinerea* (Bethke et al., 2015). In addition, Arabidopsis *response regulator 6* (*arr6*) mutant, containing a higher amount of HG, showed more resistance to *Plectosphaerella cucumerina* but more susceptibility to *Ralstonia pseudosolanacearum* (Bacete et al., 2020). Moreover, changes in HG composition and properties, such as the pattern and degree of methylesterification, have been described in different pathosystems and is one of the factors that controls plant resistance or susceptibility (Lionetti et al., 2012 and 2017; Coculo and Lionetti, 2022). The methylesterifcation degree and pattern of HG is finely modulated by pectin methylesterases (PMEs) and by the action of their respective inhibitors (PMEIs) (Levesque-Tremblay et al., 2015). These changes in HG methylation are also related to the activities of other important pectin-modifying enzymes such as polygalacturonase (PG) and pectin lyase (PL) (Levesque-Tremblay et al., 2015; Coculo and Lionetti, 2022). All these activities influence the CW biochemical properties (Levesque-Tremblay et al., 2015) such as the formation of the egg-box structure between two de-methylesterified HG domains (Mohnen, 2008), which produces stronger pectin gels that interfere with pathogen penetration, limiting the accessibility of its CW-degrading enzymes (CWDEs) (Lionetti et al., 2007; Raiola et al., 2011).

Another example of the importance of these pectin-modifying enzymes can be seen during the infection of *P. syringae* pv. *macuolicola* in Arabidopsis, which leads to an increment of PME activity (Bethke et al., 2014). In concordance with this, a PMEI3 identified in cotton enhances resistance to *Verticilium* (Liu et al., 2018). In the same way, *pme17* mutants showed an increase in susceptibility to *B. cinerea* (Corpo et al., 2020). Similarly, *pmei* mutants such as *pmei10*, *pmei11*, and *pmei12* lead to compromised immunity against *B. cinerea* (Lionetti et al., 2017). All this research confirms that pectins, more particularly the HG structure, seem to be important in the CW reinforcement and protection against Pph as well as other pathogens, but until now little is known about the role of pectin-modifying enzymes during the pathogenesis processes in common bean. On the one hand, at short times after pathogenic attack, a transcriptional reprogramming takes place, which is indicative of alterations in important cell mechanisms in order to block the action of the pathogen (Ngou et al., 2022).

However, there is less information about the triggering (or not) of transcriptomic changes during the first steps of bacterial infection, such as Pph, in susceptible varieties of common bean. On the other hand, due to the protagonist role of HG described in Arabidopsis defense against different pathogens (Bethke et al., 2014; 2015), we want to determine if, in our system, the HG domains are a key factor in common bean defense against Pph. To evaluate this, we developed transcriptional analyses and evaluated the CW changes during the first stages of bacterium colonization.

For these purposes, 15-day-old common bean plants were infected or not with Pph. Their leaves were collected at 0, 2 and 9 h post infection and their RNA and CWs were extracted for the subsequent analyses. The transcriptomic analysis has revealed the modulation of several CW-related genes implicated in pectin synthesis and modification. Among these, a common bean PMEI (PvPMEI3) protein – phylogenetically related to AtPMEI3 —could be associated with pathogenesis processes. Overexpressed at 2 hours and suppressed at 9 hours upon pathogen appearance, this protein has been proved to be located in the apoplast and could play a role in Pph infection, as Arabidopsis *pmei3* mutant showed susceptibility to Pph, in contrast to Col-0 plants. In this line, the changes in the expression profile of CW-related genes at early times post-infection could result in CW modifications, characterized by changes in pectin structure, such as in HG methylesterification and the formation of egg-box epitopes, which could be an attempt to reinforce the CW structure, and thus avoiding the attack of the bacterial CWDEs. However, these transcriptional and CW-remodeling processes are neither maintained during the necessary time, nor they block the action of the pathogen, resulting in the well-known susceptibility of *riñón* variety to Pph.

This type of studies supports the importance of pectins in defense, more particularly the participation of HG molecules, allowing the possibility to prime plants for modulating the pectin component in order to obtain crops more resistant to diseases.

## RESULTS

### Damage symptoms are quickly visible on *P. vulgaris var. riñón* leaves after Pph infection

Plant immune-signaling pathways and transcriptional reprogramming have been proved to be activated within the first 12 hours after the apparition of the pathogen (Ngou et al., 2022). Recently, De la Rubia et al. (2021a) have demonstrated that the expression of *PR1* and other plant defense-related genes increased past 2 hours upon Pph infection in *P. vulgaris var. riñón*. Therefore, we decided to perform a transcriptome analysis by RNA-seq comparing control and Pph-infected common bean leaves at 0, 2 and 9 hours post-infection (Figure 1A). To ensure that Pph had infected our plants, we performed a conventional PCR to detect the presence of Pph with specific primers for an infection marker described by Seok Cho et al. (2010), which indicated that the infection had been successful at 2 and 9 hours (Figure 1B). In parallel, a DAB staining was carried out, indicating H_2_O_2_ accumulation (ROS) due to Pph infection. This staining revealed higher H_2_O_2_ levels in infected leaves at 2 hours and, especially, at 9 hours than in control leaves, confirming that the infection had effectively taken place (Figure 1C). According to these results, we selected these two time points after Pph inoculation (2 and 9 hours) to perform an RNA-seq analysis in order to understand the molecular regulation processes that take place when Pph begins to infect and colonize common bean leaves.

**Figure 1:**
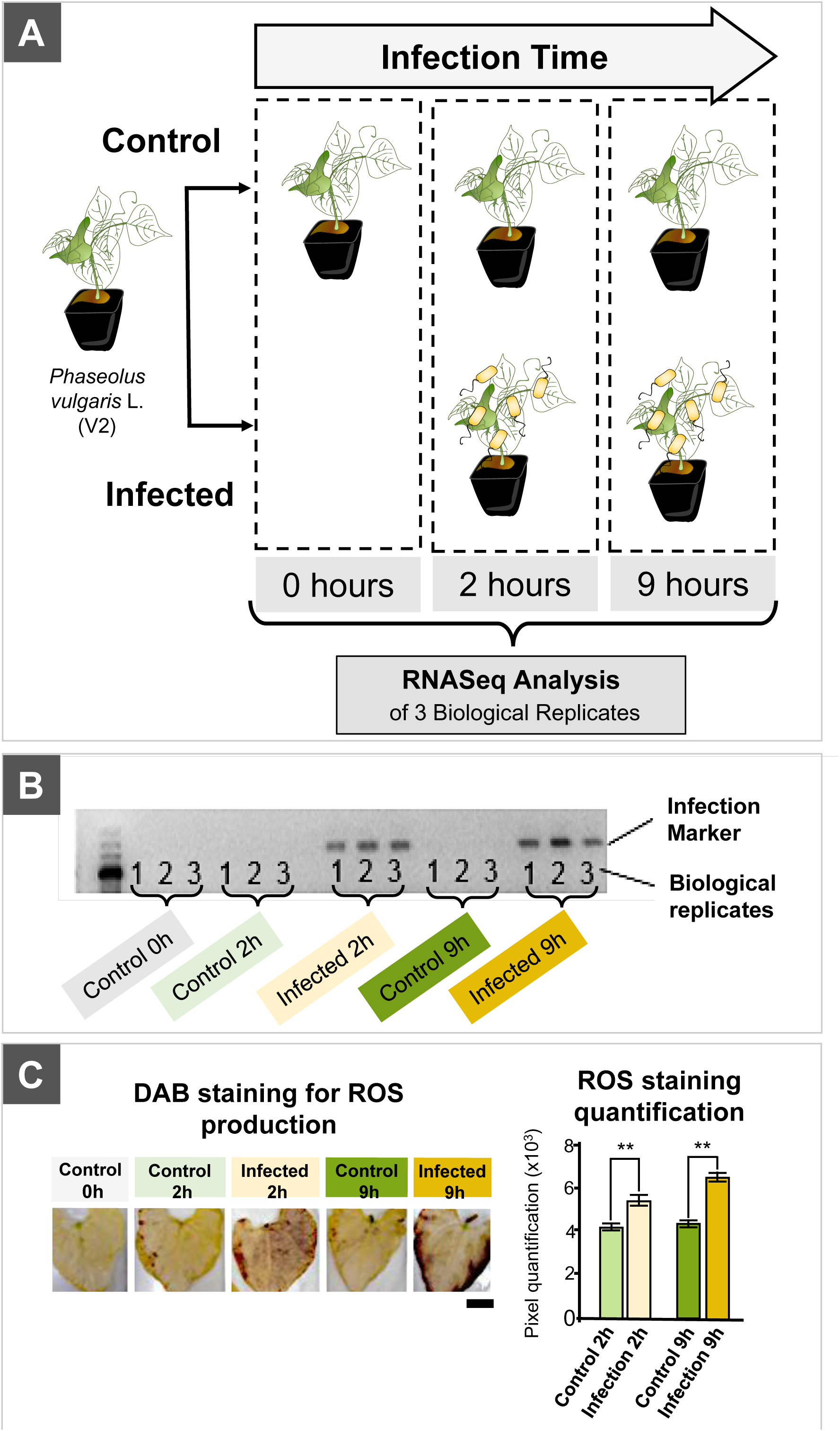
*Pseudomonas syringae* pv. *phaseolicola* (Pph) causes visible damage in common bean leaves at 2 and 9 hours post infection. *A. Phaseolus vulgaris* L. variety *riñón* plants were grown *in vitro* until V2 developmental stage. The Pph infection was caused by spraying the bacteria solution (infected plants) or water (control plants) directly on leaves. Leaves of at least 10 infected and control plants were collected at 2 and 9 hours after the treatments. Three independent experiments were performed to obtain three biological replicates. B. Expression of a Pph molecular marker (210 bp) showed the presence of the bacterium at 2 and 9 hours after the treatment only in infected leaves. C. Left panel: DAB staining of common bean leaves showed ROS-mediated damage at 2 and 9 hours after the infection with Pph. Bar: 2 cm. Right panel: The quantification of leaf damage was performed by analyzing the density of dark spots. Error bars represent the SE. Asterisks show statistical differences with one-way ANOVA, post-hoc Tukey test (p<0.05).

### Transcriptome analysis of common bean leaves revealed a high mis-regulation of several genes during the first hours of Pph infection

An RNA-seq analysis of Pph-infected and non-infected common bean leaves from the three independent infection experiments was performed, obtaining a total of 1.352.269.808 raw reads (BioProject PRJNA911667) with an average of 90.151.321 reads for each replicate, with a maximum and minimum of 122.399.890 and 68.242.664 respectively. Next, raw reads were mapped, obtaining a range of aligned reads against the *P. vulgaris* v2.1 reference genome from 89,71 % to 93,65 % (Supplementary Table 1). A Principal Component Analysis (PCA) and a heatmap were made using the 100 most differentially expressed genes (DEGs) (Figure 2A and Supplementary Figure 1), which showed that the 3 independent replicates used for the transcriptomic analysis had a treatment-related distribution with a little dispersion of the replicates due to the natural biological variability of the plants.

**Figure 2:**
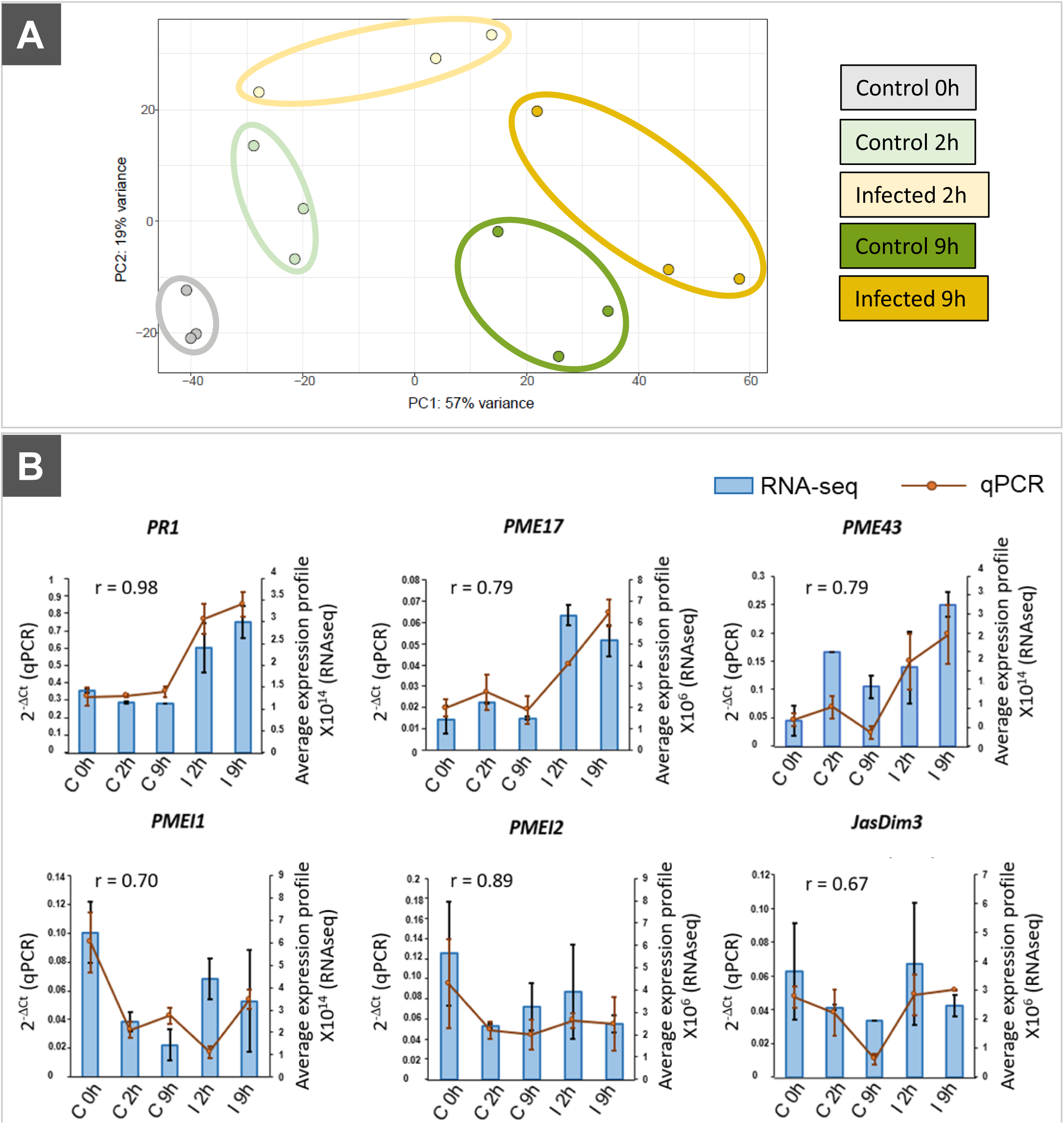
Transcriptomic analysis validation. A. Principal component analysis (PCA) using differentially expressed genes (DEGs) for each biological replicate of all Pph-infected and control samples that were used for the transcriptomic analysis. The PCA showed that, although all the experiments were independent, biological replicates had similar behavior. B. qRT-PCR validation of the RNA-seq experiment. Gene expression patterns from the RNA-seq analysis (Average expression profile, blue bars) were validated for six representative genes by qRT–PCR (2-DCt, brown line). The confirmation includes three overexpressed genes (PR1/ Phvul.003G109100, PME17/ Phvul.010G079900 and PME41/ Phvul.001G209100) and three repressed genes (PMEI1/ Phvul.002G060750, PMEI2/ Phvul.010G041500 and JasDim3/ Phvul.010G041500). The co-expression coefficient was calculated by Pearson’s test (r) using GraphPad Prism V.8. Error bars represent SE values from three biological replicates (n=9).

To validate the expression pattern observed in the transcriptomic analysis, a qRT-PCR was performed using three up-regulated (*PR1*/ Phvul.003G109100, *PME17*/ Phvul.010G079900 and *PME41*/ Phvul.001G209100) and three down-regulated (*PMEI1*/ Phvul.002G060750, *PMEI2*/ Phvul.010G041500 and *JasDim3*/ Phvul.010G041500) genes compared to control plants at 0 hours. The fold change values from these RNA-seq and qRT-PCR were represented together with the Pearson’s correlation test data, which revealed that all genes showed a similar expression trend, since the correlation value (r) was always higher than 0.67, validating the strength of the RNA-seq analysis (Figure 2B).

RNA-seq results indicate that many genes changed their expression profile due to the time factor, as a total of 437 and 3,877 genes were DEGs, mostly upregulated, in control plants at 0 hours *versus* control at 2 hours and 9 hours, respectively (Figure 3A). This fact was also reflected in the DEGs observed in infected plants at 2 and 9 hours compared to control at 0 hours, where 4,669 and 5,391 genes were deregulated, but being half up-and half down-regulated in this case (Figure 3A). These results could indicate a gene expression change due to circadian rhythms. In order to eliminate this factor, comparisons between infected and control plants at each time were made. This showed that most of the genes were deregulated at 2 hours post infection (1,575 DEGs), being mostly up-regulated (1,076), whereas the number of DEGs decreased to 565 at 9 hours post infection. This could be explained by the fact that *P. vulgaris* var. *riñón* is susceptible to Pph (Monteagudo et al., 2006; De la Rubia et al., 2021a) and the plant would trigger some early defense responses at 2 hours, which would be successfully stopped by the pathogen effectors at 9 hours post infection, diminishing the plant responses to Pph.

**Figure 3:**
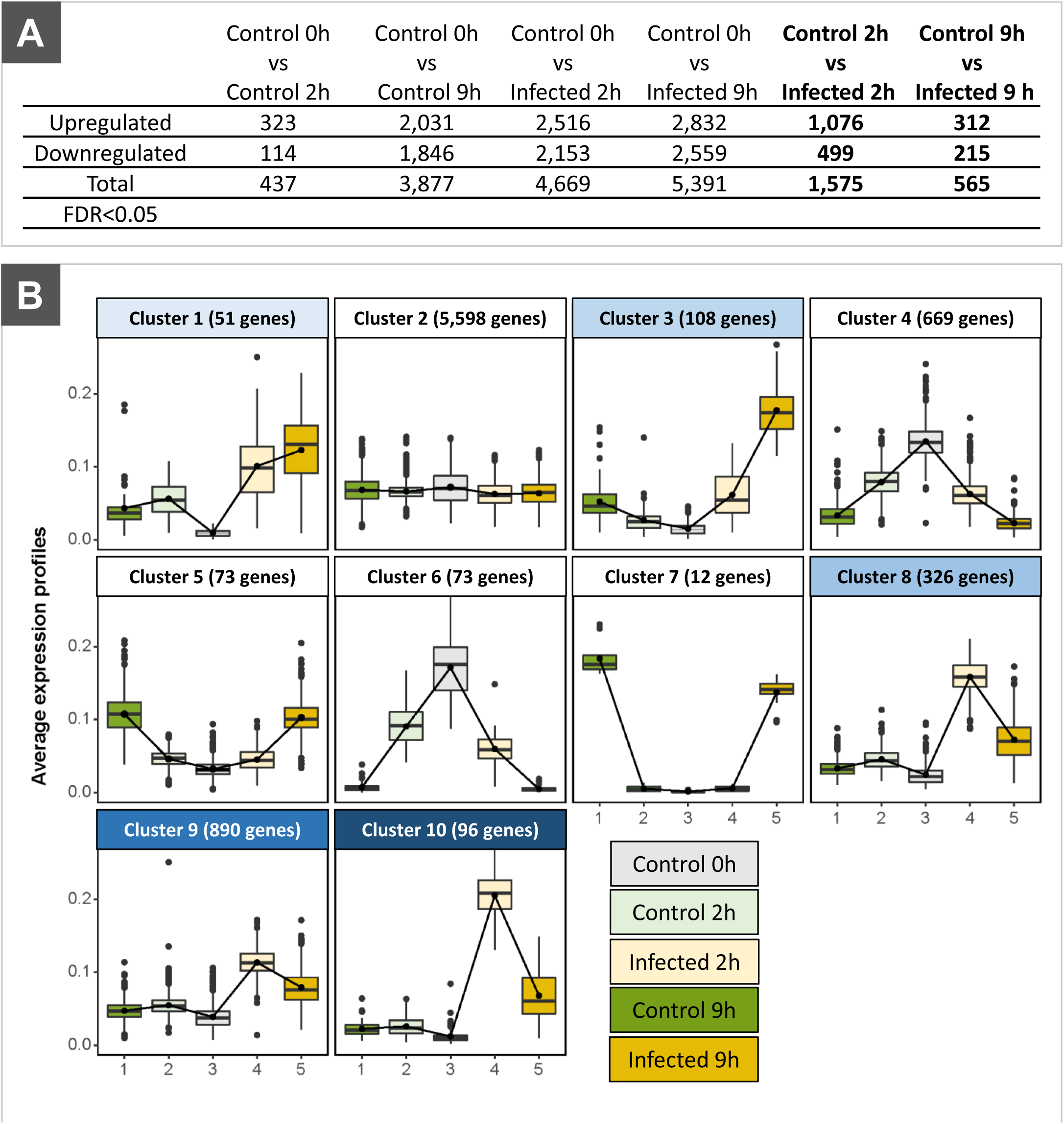
RNA-seq analysis revealed differentially expressed genes (DEGs) associated with the presence of *Pseudomonas syringae* pv *phaseolicola* (Pph) and infection time. A. Number of upregulated and downregulated genes (FDR < 0.05) due to time factor and Pph infection in infected and control plants. B. Consensus clustering based on DEGs from all Pph-infected and control plants. Clusters 1, 3, 8, 9 and 10 contained the upregulated genes associated with Pph infection throughout time, with two types of average expression profile: cluster 1 and 3 increased the upregulation of the genes with the time; cluster 8, 9 and 10 showed upregulation at 2 hours and then a slight decrease of expression at 9 hours.

Considering all DEGs, a cluster analysis was performed (Figure 3B). Consequently, the genes were classified into 10 different clusters according to their expression trend among samples. Most genes (5,598 in Cluster 2) did not show significant changes across all conditions assayed. If we focus on the selection of genes that changed their expression only due to the Pph infection, most of the DEGs belong to Cluster 8 (326 genes), Cluster 9 (890 genes) and Cluster 10 (96 genes), whose expression was increased at 2 hours and then reduced at 9 hours post infection (Figure 3B). Considering these results, most of the Pph infection responsive DEGs did not maintain their overexpression during the infection, therefore, this could be related to the susceptibility *of P. vulgaris* to Pph (Monteagudo et al., 2006; De la Rubia et al., 2021). Moreover, there were DEGs in Cluster 1 (with 51 genes) and Cluster 3 (with 108) which were mostly up-regulated at 9 hours post infection. Finally, Cluster 4 (669 genes), Cluster 5 (73 genes), Cluster 6 (73 genes) and Cluster 7 (12 genes) contained DEGs with similar profiles in control and infected plants (at 2 and 9 hours) in comparison with control at 0 hours (Figure 3B).

### Common bean responds with a transcriptomic and hormonal activation of defensive mechanisms after Pph infection

Cluster 8 and 9, which contained a higher number of genes affected by infection, were selected to perform a gene ontology (GO) analysis (Supplementary Figures 2 and 3).Interestingly, several enriched GO terms were related directly to plant defense, such as the GO terms “response to fungus”, “immune system process”, “response to wounding”, “innate immune response” or “immune response” in Cluster 8 (Supplementary Figure 2), and in Cluster 9 (Supplementary Figure 3). These results would confirm a rapid reprogramming in the transcriptional profile of common bean within hours after Pph infection. Many of the genes that were beyond these GO terms are included in Supplementary Table 2, and were described as receptors involved in the recognition of the pathogen and in initiating the defense response.

In addition, other GO terms were related to infection processes, such as those associated with “response to salicylic acid” only in Cluster 8 (Supplementary Figure 2) and “response to jasmonic acid” in both Cluster 8 and 9 (Supplementary Figure 3) and the misregulated genes associated with these PHs appear in Supplementary Figure 4. SA and JA, together with ethylene and abscisic acid (ABA), are the major signaling regulators in plant defense (Anver and Tsuda, 2015; Gupta et al., 2017). For this reason, the concentration of these PHs was also measured. Although a quick accumulation of SA and JA was observed at 2 hours post infection, the concentrations of these two PHs decreased to non-infected levels at 9 hours (Figure 4.A and C). In the case of ABA and MeJA, no differences were found at the two time points tested after infection (Figure 4.B and D). Those results suggest that, although common bean plants attempted to trigger defense responses, such responses did not endure in time long enough to defend the plant against the pathogen efficiently.

**Figure 4:**
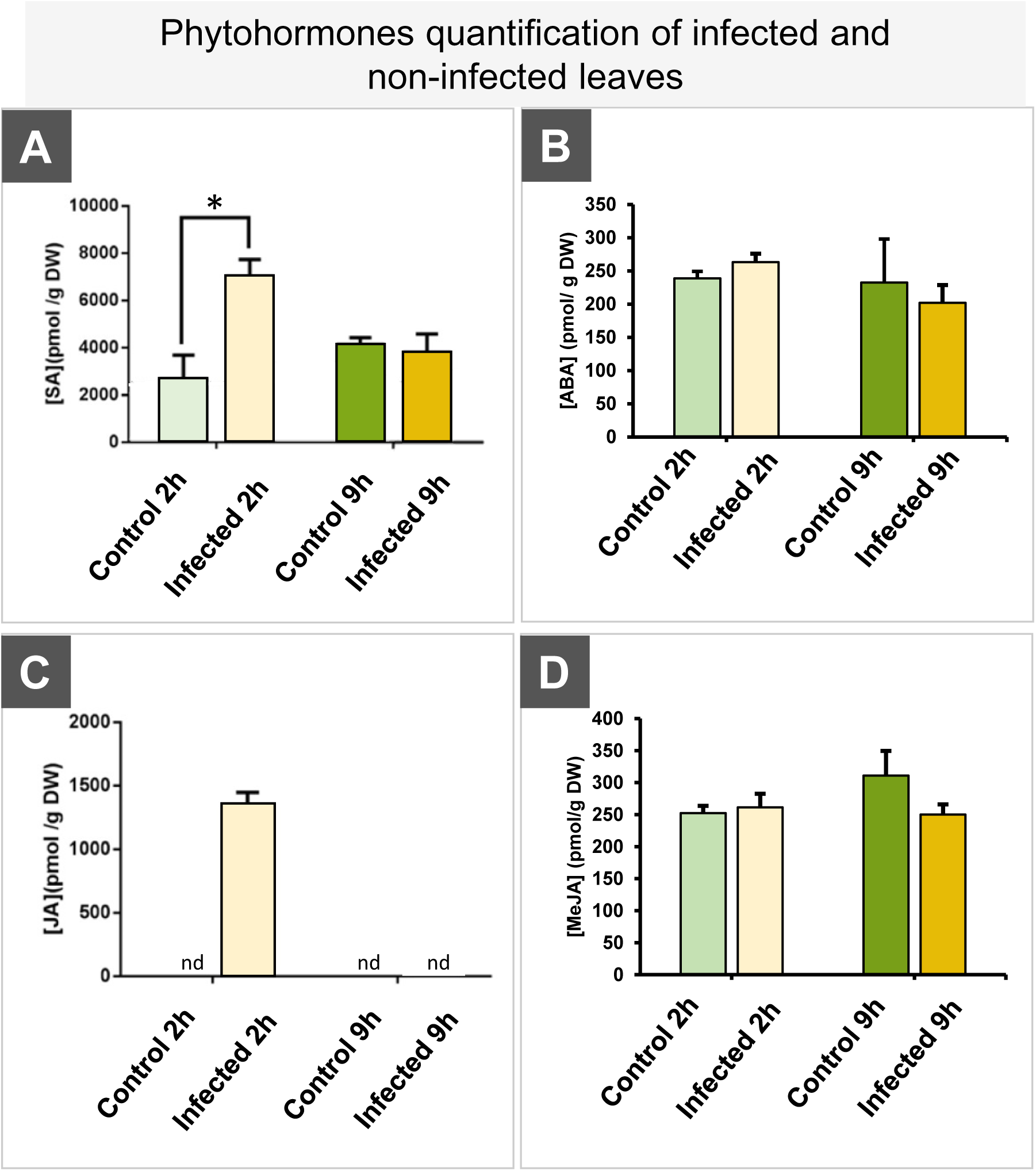
*Pseudomonas syringae* pv *phaseolicola* (Pph) infection of common bean leaves showed a quick Salicilic Acid (SA) and Jasmonic Acid (JA) accumulation at 2 hours post infection, which decreased to normal levels at 9 hours. SA (A), ABA (B), JA (C) and MeJA (D) content in infected and non-infected common bean leaves at 2 and 9 hours after inoculation. Error bars show the SE from three biological replicates. Asterisks represent significant differences (p<0.05) using Student’s t-test.

### Pph infection provoked the overexpression of a high number of CW-related genes

The transcriptomic analysis revealed a high number of DEGs associated with CW metabolism in all the clusters related to Pph-infection (Table 1). A total of 101 CW-related DEGs were detected, most of them belonging to clusters 8 and 9, in which overexpression was observed at 2 hours after infection (Figure 3, Table 1). Interestingly, from all CW-related DEGs, the genes involved in polysaccharide biosynthesis (mainly putative glycosyltransferases) and in CW remodeling (mainly putative glycosyl hydrolases) were the two groups with the highest number of genes overexpressed (36 and 23, respectively). These results suggest that the Pph infection should promote a quick and extensive CW remodeling. In addition, there were also 9 genes related to pectin metabolism which were overexpressed at 2 hours after Pph infection (Table 1). Moreover, lignin (18 genes) and callose (6 genes) biosynthesis genes were also overexpressed (Table 1). The synthesis of both molecules has been associated with plant defense responses to biotic stresses (Caño-Delgado et al., 2003; Miedes et al., 2014; Wang et al., 2021;). Finally, CW-protein genes, such as expansins (3 genes) and glycoproteins (3 genes) (Table 1) were also identified.

**Table 1.** Cell wall-related DEGs belonging to clusters selected in the transcriptomic analysis of the first steps of *Pseudomonas syringae* pv. *phaseolicola* (Pph) infection of common bean leaves. The metabolic process, gene ID, cluster number, symbols and gene description of each gene are shown in the table.

|  | GeneID | Cluster | Symbols | Description |
| --- | --- | --- | --- | --- |
| Cell wall polysaccharide biosynthesis | Phvul.008G175500.1 | 1 | UGT85A2 | UDP-glucosyl transferase 85A2 |
|  | Phvul.005G116501.1 | 3 | CSLB03 | Cellulose synthase-like B3 |
|  | Phvul.005G117833.1 | 3 | CSLB04 | Cellulose synthase-like B4 |
|  | Phvul.001G183000.1 | 3 | UGT73D1 | UDP-glucosyl transferase 73D1 |
|  | Phvul.005G005400.1 | 3 | UGT88A1 | UDP-glucosyl transferase 88A1 |
|  | Phvul.002G168400.1 | 3 |  | Glycosyl transferase, family 35 |
|  | Phvul.011G138600.1 | 3 |  | UDP-Glycosyltransferase superfamily protein |
|  | Phvul.006G058700.1 | 8 | CSLE1 | Cellulose synthase like E1 |
|  | Phvul.008G140500.1 | 8 | UGT72B1 | UDP-Glycosyltransferase superfamily protein |
|  | Phvul.007G251200.1 | 8 |  | UDP-Glycosyltransferase superfamily protein |
|  | Phvul.008G034000.1 | 8 |  | UDP-Glycosyltransferase superfamily protein |
|  | Phvul.008G140500.1 | 8 | UGT72B1 | UDP-Glycosyltransferase superfamily protein |
|  | Phvul.008G140600.1 | 8 | UGT72B1 | UDP-Glycosyltransferase superfamily protein |
|  | Phvul.011G080500.1 | 8 |  | Glycosyltransferase family protein 2 |
|  | Phvul.003G146700.1 | 8 |  | Galactosyltransferase family protein |
|  | Phvul.002G136300.1 | 9 | CSLD3 | cellulose synthase-like D3 |
|  | Phvul.006G058600.1 | 9 | CSLE1 | cellulose synthase like E1 |
|  | Phvul.008G140800.1 | 9 | UGT72B1 | UDP-Glycosyltransferase superfamily protei |
|  | Phvul.003G277200.1 | 9 |  | UDP-Glycosyltransferase superfamily protein |
|  | Phvul.006G205600.1 | 9 | UGT85A1 | UDP-Glycosyltransferase superfamily protein |
|  | Phvul.007G039200.1 | 9 | UGT84A1 | UDP-Glycosyltransferase superfamily protein |
|  | Phvul.008G257200.1 | 9 |  | UDP-Glycosyltransferase superfamily protein |
|  | Phvul.008G140700.1 | 9 | UGT72B1 | UDP-Glycosyltransferase superfamily protein |
|  | Phvul.004G103900.1 | 9 | UGT88A1 | UDP-glucosyl transferase 88A1 |
|  | Phvul.002G026900.1 | 9 | UGT72E1 | UDP-glucosyl transferase 72E1 |
|  | Phvul.003G047200.2 | 9 | UGT73B4 | UDP-glycosyltransferase 73B4 |
|  | Phvul.007G039600.1 | 9 | UGT84B1 | UDP-glucosyl transferase 84B1 |
|  | Phvul.008G174700.1 | 9 | UGT85A2 | UDP-glucosyl transferase 85A2 |
|  | Phvul.011G060300.1 | 9 | UGT76E11 | UDP-glucosyl transferase 76E11 |
|  | Phvul.011G151200.1 | 9 | UGT89B1 | UDP-glucosyl transferase 89B1 |
|  | Phvul.008G225700.1 | 9 | EPC1 | Nucleotide-diphospho-sugar transferases superfamily protein |
|  | Phvul.002G089000.1 | 9 | UTR3 | UDP-galactose transporter 3 |
|  | Phvul.002G084400.1 | 9 | GALT1 | Galactosyltransferase 1 |
|  | Phvul.003G251200.1 | 9 | GATL2 | Galacturonosyltransferase-like 2 |
|  | Phvul.008G035400.1 | 9 | GAUT13 | Galacturonosyltransferase 13 |
|  | Phvul.009G026100.1 | 9 |  | Galactosyltransferase family protein |
| Cell wall remodelling | Phvul.011G055900.1 | 1 | BGLU15 | β--glucosidase 15 |
|  | Phvul.011G077900.1 | 3 |  | Glycosyl hydrolase superfamily protein |
|  | Phvul.011G177000.1 | 3 |  | Glycosyl hydrolase superfamily protein |
| | Phvul.011G056100.1 | 8 | BGLU13 | β--glucosidase 13 |
| | Phvul.001G076700.1 | 8 | BGLU17 | β--glucosidase 17 |
| | Phvul.011G055800.1 | 8 | BGLU17 | β--glucosidase 17 |
|  | Phvul.010G137800.1 | 8 | BFRUCT1 | Glycosyl hydrolases family 32 protein |
|  | Phvul.008G033000.1 | 8 |  | Glycosyl hydrolase family 81 protein |
|  | Phvul.001G046400.1 | 8 |  | Glycosyl hydrolase family protein with chitinase insertion domain |
| | Phvul.001G128500.1 | 9 | BG1 | β-1,3-glucanase 1 |
| | Phvul.001G128550.1 | 9 | BG1 | β--1,3-glucanase 1 |
| | Phvul.008G097700.1 | 9 | BGLU11 | β-glucosidase 11 |
| | Phvul.009G127500.1 | 9 | BGAL16 | β-galactosidase 16 |
| | Phvul.003G075400.1 | 9 | | β-glucosidase, GBA2 type family protein |
|  | Phvul.001G112700.1 | 9 |  | Glycosyl hydrolase family protein |
|  | Phvul.007G250300.1 | 9 |  | Glycosyl hydrolase family protein |
|  | Phvul.011G176950.1 | 9 |  | Glycosyl hydrolase family protein |
|  | Phvul.006G092100.1 | 9 |  | Glycoside hydrolase family 2 protein |
|  | Phvul.001G123000.2 | 9 | RSW3 | Glycosyl hydrolases family 31 protein |
|  | Phvul.005G158500.1 | 9 | BFRUCT1 | Glycosyl hydrolases family 32 protein |
|  | Phvul.004G049200.1 | 9 |  | Glycosyl hydrolase family 81 protein |
|  | Phvul.008G033100.1 | 9 |  | Glycosyl hydrolase family 81 protein |
|  | Phvul.008G033200.1 | 9 |  | Glycosyl hydrolase family 81 protein |
| Pectin metabolism | Phvul.001G209100.1 | 8 | PME41 | Pectin methylesterase superfamily protein |
|  | Phvul.003G217200.1 | 8 | PMEI3 | Pectin methylesterase inhibitor family protein |
|  | Phvul.003G227500.1 | 8 | PMEI | Pectin methylesterase inhibitor family protein |
|  | Phvul.001G140000.1 | 8 | PGIP1 | Polygalacturonase inhibiting protein 1 |
|  | Phvul.006G071800.1 | 9 | PMEI | Pectin methylesterase inhibitor family protein |
|  | Phvul.008G183500.1 | 9 |  | Rhamnogalacturonate lyase family protein |
|  | Phvul.010G079900.1 | 9 | PME17 | Pectin methylesterase superfamily |
|  | Phvul.002G201800.1 | 9 | PGIP1 | Polygalacturonase inhibiting protein 1 |
|  | Phvul.002G201900.1 | 9 | PGIP1 | Polygalacturonase inhibiting protein 1 |
| Expansins | Phvul.009G186400.1 | 1 | EXP15 | Expansin A15 |
|  | Phvul.003G228000.1 | 8 | EXLB1 | Expansin-like B1 |
|  | Phvul.003G224800.1 | 9 | EXLB1 | Expansin-like B1 |
| Glycoproteins | Phvul.004G098000.1 | 3 | EXT3 | Extensin 3 |
|  | Phvul.007G063100.1 | 8 |  | Hydroxyproline-rich glycoprotein family protein |
|  | Phvul.002G158200.1 | 9 |  | Hydroxyproline-rich glycoprotein family protein |
| Callose | Phvul.002G160700.1 | 9 | RBOHF | Respiratory burst oxidase protein F |
|  | Phvul.005G162300.1 | 9 |  | Leucine-rich repeat protein kinase family protein |
|  | Phvul.005G166500.1 | 9 | PDR8 | Plant PDR ABC-type transporter family protein |
|  | Phvul.008G252300.1 | 9 | EXO70B1 | Exocyst subunit exo70 family protein B1 |
|  | Phvul.009G062400.1 | 9 | MAPKKK5 | Mitogen-activated protein kinase souvenir kinase 5 |
|  | Phvul.002G196200.1 | 9 |  | LRR-like protein kinase family protein |
| Lignin | Phvul.010G120200.1 | 3 | OMT | O-methyltransferase |
|  | Phvul.007G091000.1 | 3 | OMT | O-methyltransferase |
|  | Phvul.001G177700.2 | 8 | PAL | PHE ammonia lyase |
|  | Phvul.001G177800.1 | 8 | PAL | PHE ammonia lyase |
|  | Phvul.007G150500.1 | 8 | PAL | PHE ammonia lyase |
|  | Phvul.008G289500.1 | 8 | PAL | PHE ammonia lyase |
|  | Phvul.001G117900.1 | 8 | C3H | Coumarate-3-hydroxylase |
|  | Phvul.008G247400.1 | 8 | C4H | Cinnamate-4-hydroxylase |
|  | Phvul.002G040100.1 | 8 | 4CL3 | 4-coumarate:CoA ligase 3 |
|  | Phvul.003G047900.1 | 8 | CCoAOMT | Caffeoyl-coA O-methyltransferase |
|  | Phvul.006G079700.1 | 9 | C4H | Cinnamate-4-hydroxylase |
|  | Phvul.007G026000.1 | 9 | C4H | Cinnamate-4-hydroxylase |
|  | Phvul.005G184500.1 | 9 | 4CL2 | 4-coumarate:CoA ligase 2 |
|  | Phvul.009G044400.1 | 9 | OMT | O-methyltransferase |
|  | Phvul.004G172600.2 | 9 | CCoAOMT | Probable Caffeoyl-CoA O-methyltransferase |
|  | Phvul.010G160700.1 | 9 | CCR1 | Cinnamoyl coa reductase 1 |
|  | Phvul.003G006000.2 | 10 | OMT | O-methyltransferase |
|  | Phvul.002G017500.1 | 10 | CCoAOMT | Caffeoyl-coA O-methyltransferase |

### Pph infection provoked changes in the metabolism of homogalacturonan of common bean

In order to explore if the change in the expression of CW genes was also reflected in structural changes, the monosaccharide composition of the Alcohol Insoluble Residue (AIR) from infected and control plants was quantified (Figure 5A). As a result, no differences were observed at 2 hours post infection. However, an increase in arabinose and a decrease in xylose were found at 9 hours of infection. These results suggest that Pph infection causes changes in matrix polysaccharides, probably hemicelluloses (xyloglucan or xylans), and/or the presence of arabinose or xylose in RG-I and RG-II side chains. However, considering that most of pectin-related misregulated genes due to infection were associated with HG metabolism (Table 1), we quantified and compared the methanol content in Pph-infected and control leaves (Figure 5B). Methanol released from the AIR fraction tended to increase at 2 hours post infection, but the difference with respect to control was only significant at 9 hours, indicating that the HG pectic domains of infected leaves were more methylated.

**Figure 5:**
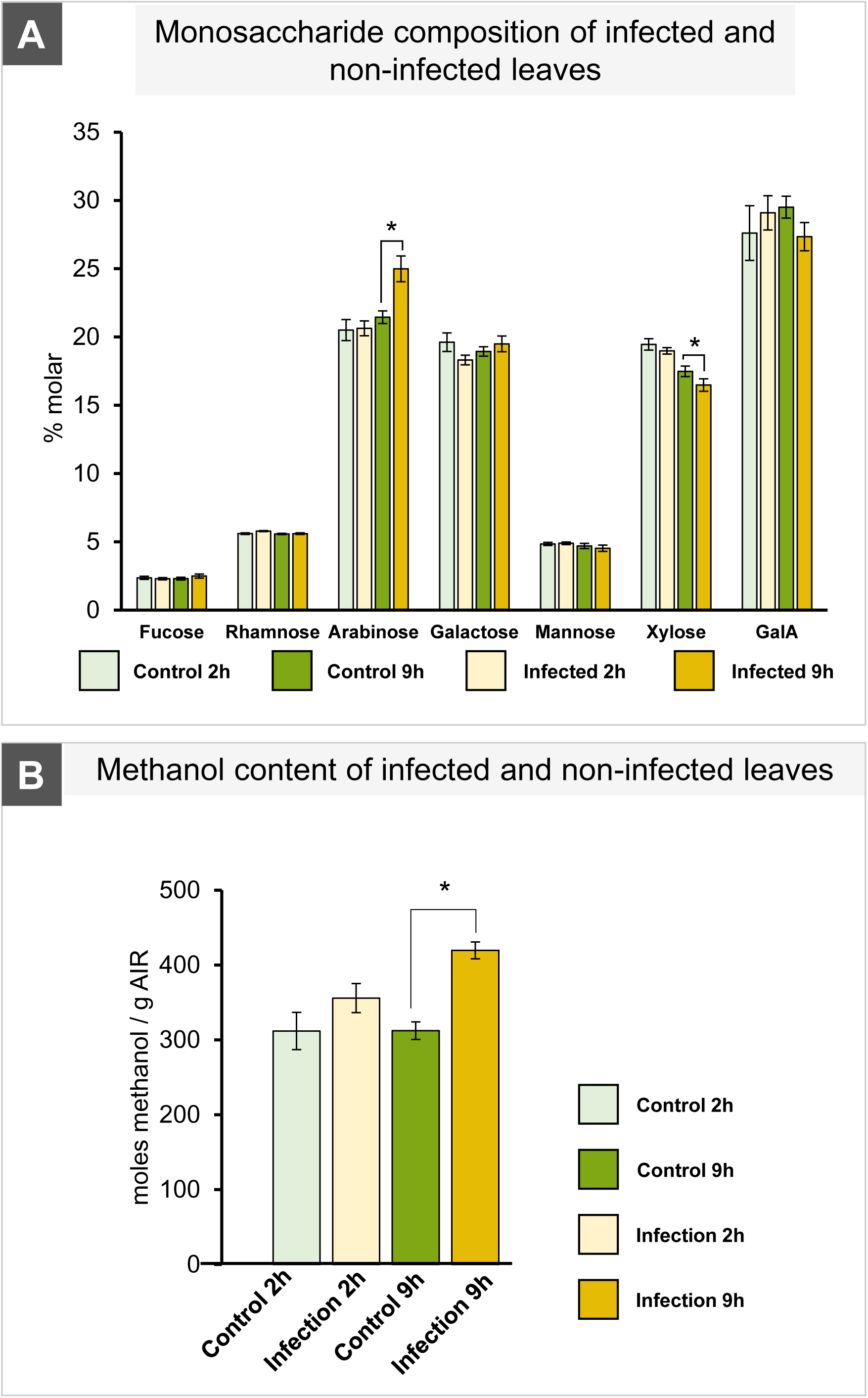
*Pseudomonas syringae* pv *phaseolicola* (Pph) infection causes changes in cell wall monosaccharide composition and polysaccharide methylesterification of common bean leaves. A. Monosaccharide composition in molar % of common bean leaves at 2 and 9 hours after Pph infection. Cell wall-derived sugar quantification was performed in alcohol insoluble residue from infected and non-infected common bean leaves using HPEAD-PAD. Error bars show the SE from three biological replicates. Asterisks represent significant differences using one-way ANOVA, Tukey test (p<0.05). B. Methanol content of infected and non-infected common bean leaves. Error bars show the SE from three biological replicates. Asterisks show significant differences using Student’s t-test (p<0.05).

To confirm if this increase in methanol content was related to changes in HG methylation, immunolabelling of leaf sections was carried out by using JIM5, JIM7 and 2F4 antibodies, which recognize partially methylesterified HG, highly methylesterified HG, and egg-box structures, respectively (Figure 6, 7 and 8). The results showed an increase of partially methylesterified HG epitopes (yellow) upon infection mainly localized in the parenchymatic tissues, especially in the palisade parenchyma (Figure 6A and 6B). The quantification of JIM5 labeling showed almost twice more signal in Pph-infected leaves than in control leaves at 2 and 9 hours post infection (Figure 6C). For JIM7 (blue), the increment in labeling of highly methylesterified HG epitopes was observed at both times of infection and this increment was detected in all the leaf tissues but was especially notable in parenchymatic cells (Figure 7A and 7B). As for JIM5, an increase in the antibody signal in response to Pph infection was measured (Figure 7C). The 2F4 antibody labeling, which targets HG dimerization by egg-box complexes, was also evaluated in Figure 8. 2F4 labeling (green) showed the detection of more egg-box complex epitopes upon infection and, interestingly, it was higher at 9 hours post infection (Figure 8.B). All these results point to important modifications in the HG structure, confirming changes in the pattern of methylesterification of HG domains, which also affected the increment in the accumulation of egg-boxes in the CW of parenchymatic tissues in response to Pph infection.

**Figure 6:**
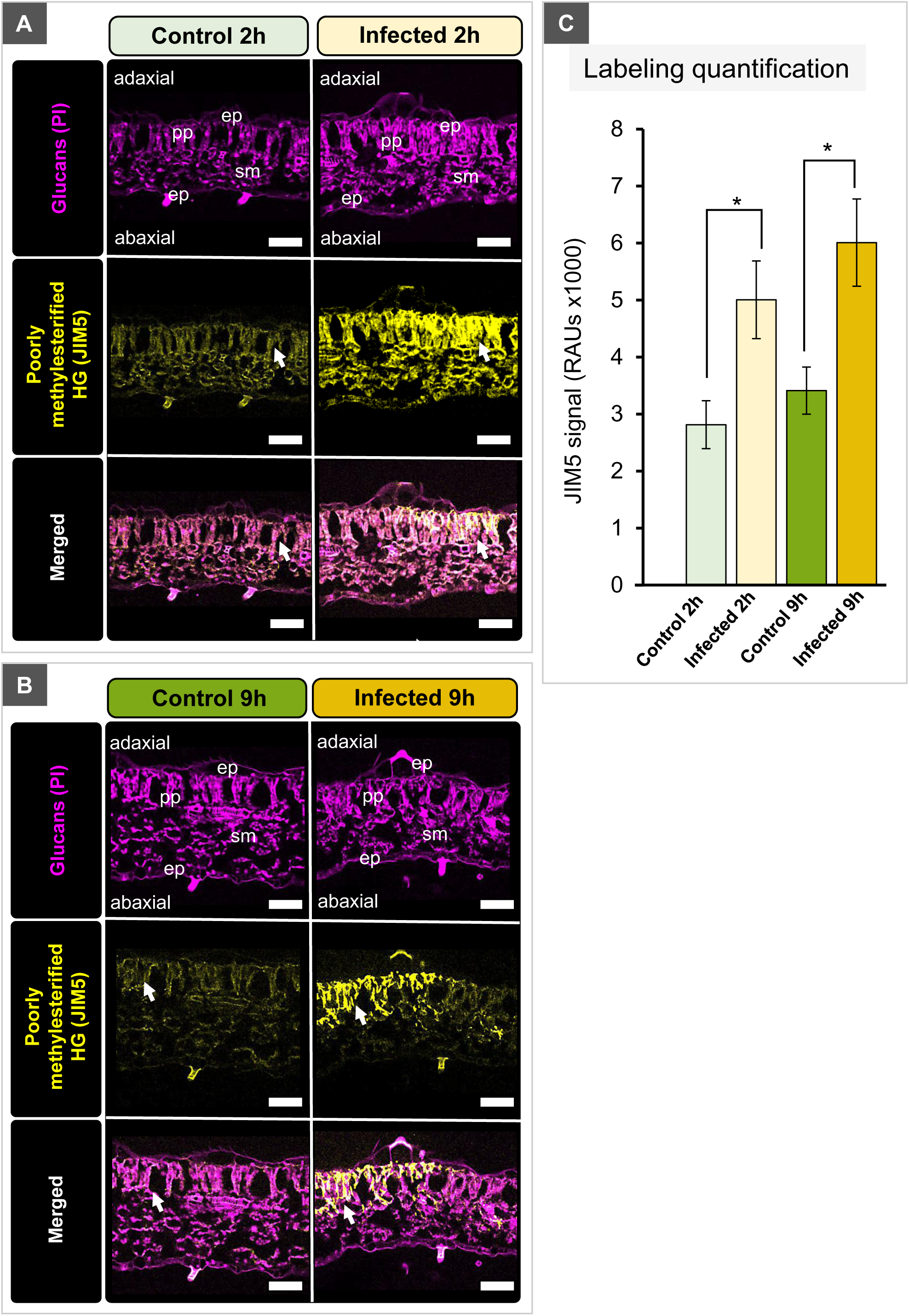
*Pseudomonas syringae* pv *phaseolicola* (Pph) infection of common bean leaves showed an accumulation of partially methylesterified homogalacturonans (HGs) mainly in the mesophyll palisade parenchyma (pp) at early times post infection. A and B. Partially methylesterified homogalacturonan using JIM5 immunolabeling in infected and non-infected common bean leaves at 2 (A) and 9 (B) hours. Reconstruction of optical sections from confocal microscopy using JIM5 (yellow) and propidium iodide to label cell walls (purple). Arrows show an accumulation of JIM5 immunolabelling in infected leaves, ep: epidermis; pp: palisade parenchyma; sm: spongy mesophyll. Bars: 50 µm C. Quantification of the JIM5 immunolabeling (yellow) from infected and non-infected leaves at 2 and 9 hours post infection. The pixel intensity was quantified in several pictures from at least three biological replicates. Asterisks represent significant differences using Student’s t-test (*p*<0.05).

**Figure 7.**
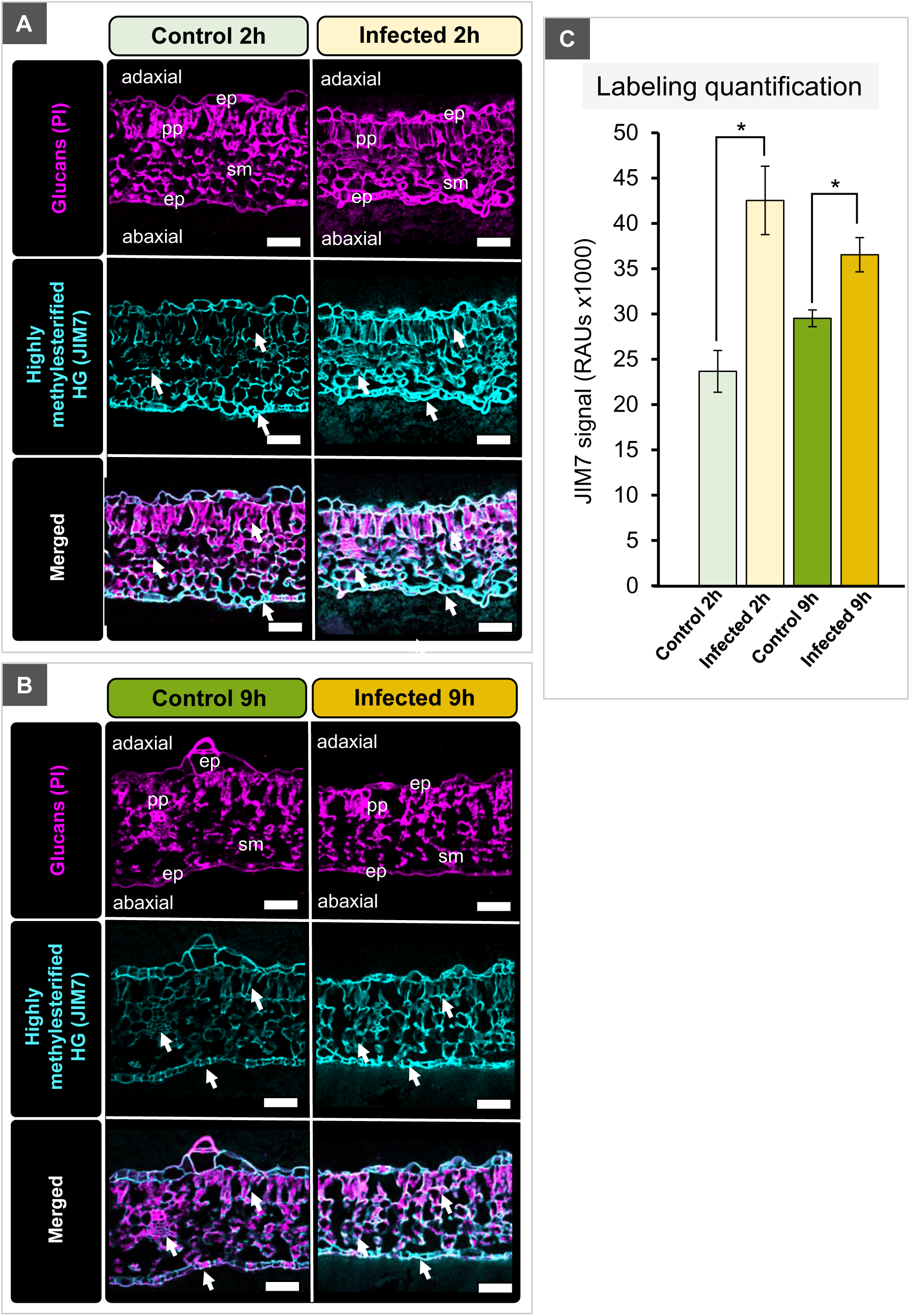
*Pseudomonas syringae* pv *phaseolicola* (Pph) infection produced a general increase in the accumulation of highly methylesterified homogalacturonans (HGs) mainly in the mesophyl of common bean leaves. A and B. Highly methylesterified homogalacturonan using JIM7 immunolabeling in infected and non-infected common bean leaves at 2 (A) and 9 (B) hours. Reconstruction of confocal microscopy optical sections using JIM7 (cyan) and propidium iodide to label cell walls (purple). Arrows show an accumulation of JIM7 immunolabelling in infected leaves, ep: epidermis; pp: palisade parenchyma; sm: spongy mesophyll. Bars: 50 µm. C. Quantification of JIM7 immunolabeling (cyan) from infected and non-infected leaves at 2 and 9 hours post infection. The pixel intensity was quantified in several pictures from at least three biological replicates. Asterisks represent significant differences using Student’s t-test (p<0.05).

**Figure 8.**
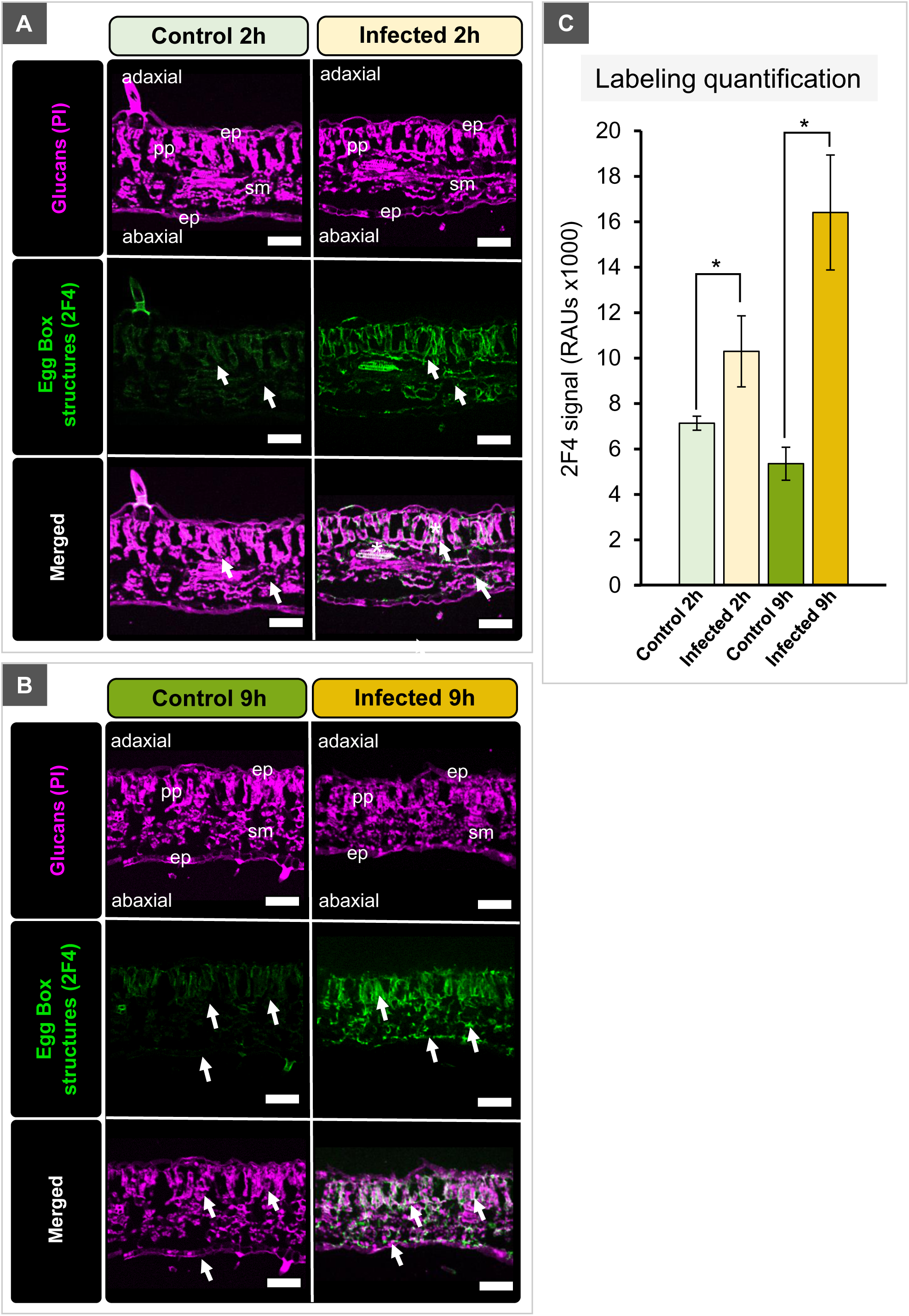
*Pseudomonas syringae* pv *phaseolicola* (Pph) infection of common bean leaves provoked the formation of dimeric associations of homogalacturonan (HG) molecules through calcium egg-boxes mainly in parenchymatic tissues. A and B. HG egg-box formation using 2F4 immunolabeling in infected and non-infected bean leaves at 2 (A) and 9 (B) hours. Reconstruction of confocal microscopy optical sections using 2F4 (green) and propidium iodide to label cell walls (purple). Arrows show an accumulation of 2F4 immunolabelling in infected leaves, ep: epidermis; pp: palisade parenchyma; sm: spongy mesophyll. Bars: 50 µm. C. Quantification of the 2F4 immunolabeling (green) from infected and non-infected leaves at 2 and 9 hours. The pixel intensity was quantified in several pictures from at least three biological replicates. Asterisks represent significant differences using Student’s t-test (*p*<0.05).

### Pectin-related enzymatic activities that protect HG from degradation are inhibited during early Pph infection

Pph infection provoked changes in HG methylesterification and the distribution of low and high-methylesterified HG populations in common bean leaves. HG modifying enzymes, such as PME, polygalacturonase (PG) and pectin lyase (PL) activities were assayed in infected and non-infected leaf extracts (Figure 9) to confirm changes in HG metabolism. Our results indicated that PME activity significantly decreased at 2 hours in Pph-infected leaves compared to control and then, at 9 hours after infection, no differences in PME activity compared to control were observed (Figure 9B). In concordance with this, PL and PG activities tended to decrease at all times post infection compared to their respective control at each time, reaching significant differences in PL activity only at 9 hours post infection (Figure 9D).

**Figure 9:**
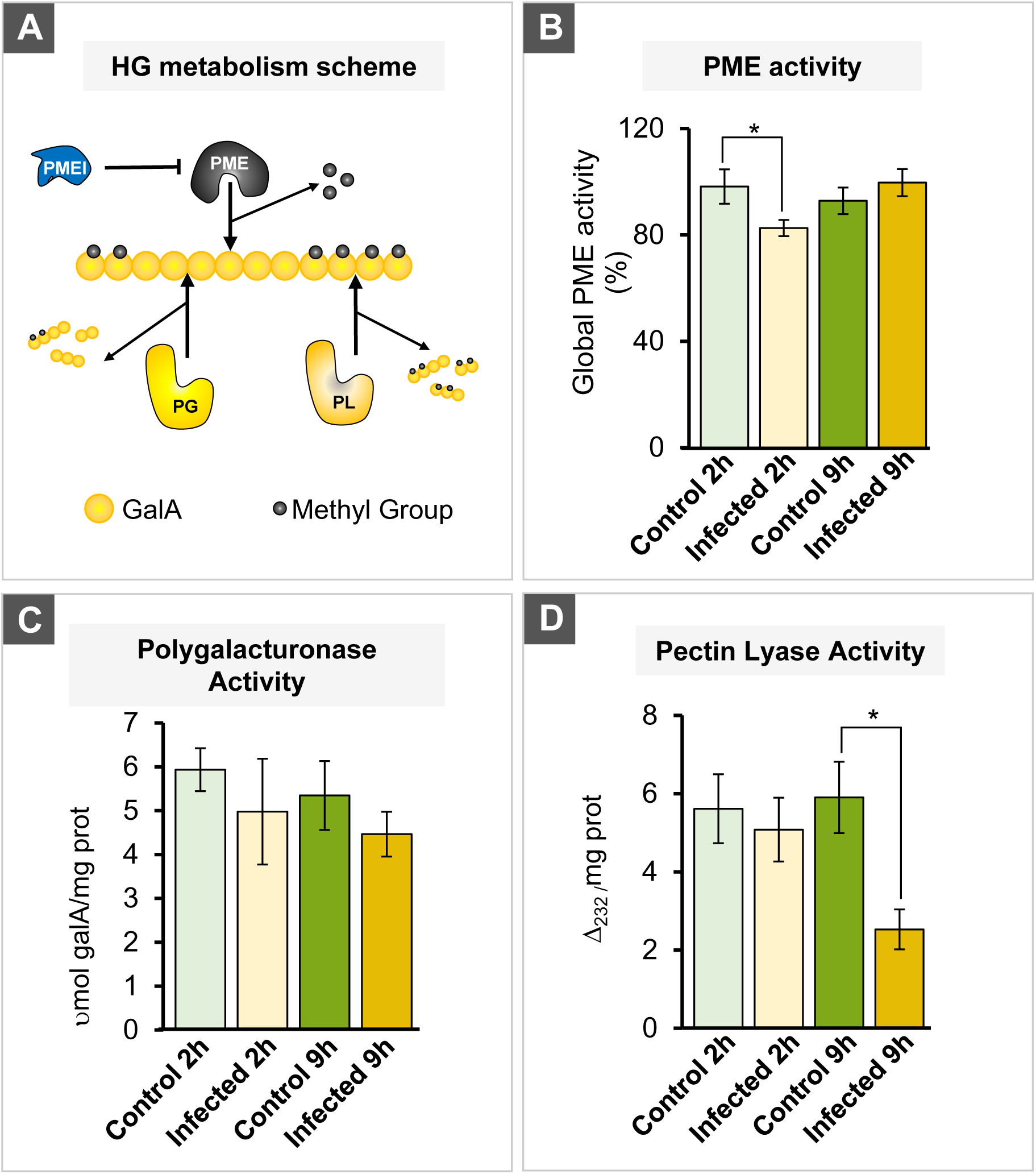
*Pseudomonas syringae* pv *phaseolicola* (Pph) infection of common bean cells caused a quick reduction of homogalacturonan (HG) metabolic enzyme activities. A. Schematic representation of the action of HG metabolic enzymes: pectin methylesterase (PME), Polygalacturonase (PGs) and Pectin Lyase (PL). PME catalyzed HG demethylesterification, and this activity is controlled by protein inhibitors (PMEIs). PGs and PLs can degrade HG molecules in different ways depending on the pattern and degree of methylesterification of HG. B. Global PME activity in infected and non-infected common bean leaves at 2 and 9 hours in three biological replicates (n=9). Asterisks represent significant differences using Student’s t-test (p<0.05). C. PG activity in infected and non-infected common bean leaves at 2 and 9 hours in three biological replicates (n=9). D. PL activity in infected and non-infected common bean leaves at 2 and 9 hours in three biological replicates (n=9). Asterisks represent significant differences using Student’s t-test (p<0.05).

These results would explain the increased methanol content detected in the CW of the leaves (Figure 5B). To test that Pph was able to produce these CWDEs, enzymatic assays were also performed in Pph grown in KB medium with glycerol (KB) and with polygalacturonic acid (PGA) supplementation (Supplementary Figure 5). These activities were assayed in the supernatant (media) and the pellet (bacteria) after a centrifugation. No PME activity was observed in any of the samples assayed (data not shown). In addition, no PME inhibitory effect on a commercial PME was found (Supplementary Figure 5C). However, results suggest that Pph secreted PL and PG into the media in the presence of PGA (Supplementary Figure 5A and 5B).

### Pectin and HG-metabolism genes were overexpressed under Pph infection in common bean leaves

As demonstrated above, the Pph infection of common bean leaves caused changes in HG domains, provoked by an alteration of the HG-specific enzyme activities. Of all the CW-related DEGs, 9 belonging in a group with a putative role in pectin metabolism were grouped in cluster 8 and 9 (overexpressed in Pph-infected plants at 2 hours and with levels similar to those of non-infected plants at 9 hours). Concretely, 3 of them would be polygalacturonase inhibiting proteins (Phvul.001G140000.1, Phvul.002G201800.1 and Phvul.002G201900.1), 1 rhamnogalacturonate lyase protein (Phvul.008G183500.1), 2 PMEs (Phvul.001G209100.1 and Phvul.010G079900.1) and 3 PMEIs (Phvul.003G217200.1, Phvul.003G227500.1 and Phvul.006G071800.1) (Table 1).

A phylogenetic analysis of all PME and PMEI proteins of Arabidopsis (AT) and *P. vulgaris* (Phvul) is displayed in Figure 10A. The tree showed three major clades: PMEs (green clade), PMEIs (yellow clade) and unclassified (blue clade) (Chen et al., 2018). Common bean PMEs Phvul.010G079900.1 and Phvul.010G079900.1 are the ortholog genes of At*PME17* and At*PME41*, which were previously described to have a role in Arabidopsis defense responses to *Botrytis* and *Pseudomonas* infection, respectively (Bethke et al., 2014; Corpo et al., 2020). Moreover, common bean PMEIs Phvul.006G071800.1 and Phvul.003G227500.1 were closely related to the previously described AtPMEI12, which has a role in Arabidopsis-*Botrytis* interaction (Figure 10A; Lionetti et al., 2017). Finally, the novel common bean PMEI, Phvul.003G217200.1 (henceforth PvPMEI3), is the ortholog to AtPMEI3 (Figure 10B, Supplementary Figure 6), which is a PMEI with a role in Arabidopsis phyllotaxis, but without a demonstrated role in plant-pathogen interactions (Peaucelle et al., 2008). PvPMEI3 was the most overexpressed PMEI in this RNA-seq study (Supplementary Figure 6) and had a similar structure in the four α-helices to that of AtPMEI3 (Figure 10B), characteristic of PMEIs: four long α-helices arranged in an up-down-up-down topology, rendering a four-helix bundle (Giovane et al., 2004; Di Matteo et al., 2005) and an N-terminal region composed of three short, distorted helices called α-hairpin module, which is important for PMEI activity (Hothorn et al 2004b). Specifically, they have a percentage of identity of 38.07 % and a query coverage of 83 % (Supplementary Figure 7). These results encouraged us to study whether this gene could participate in the infection.

**Figure 10:**
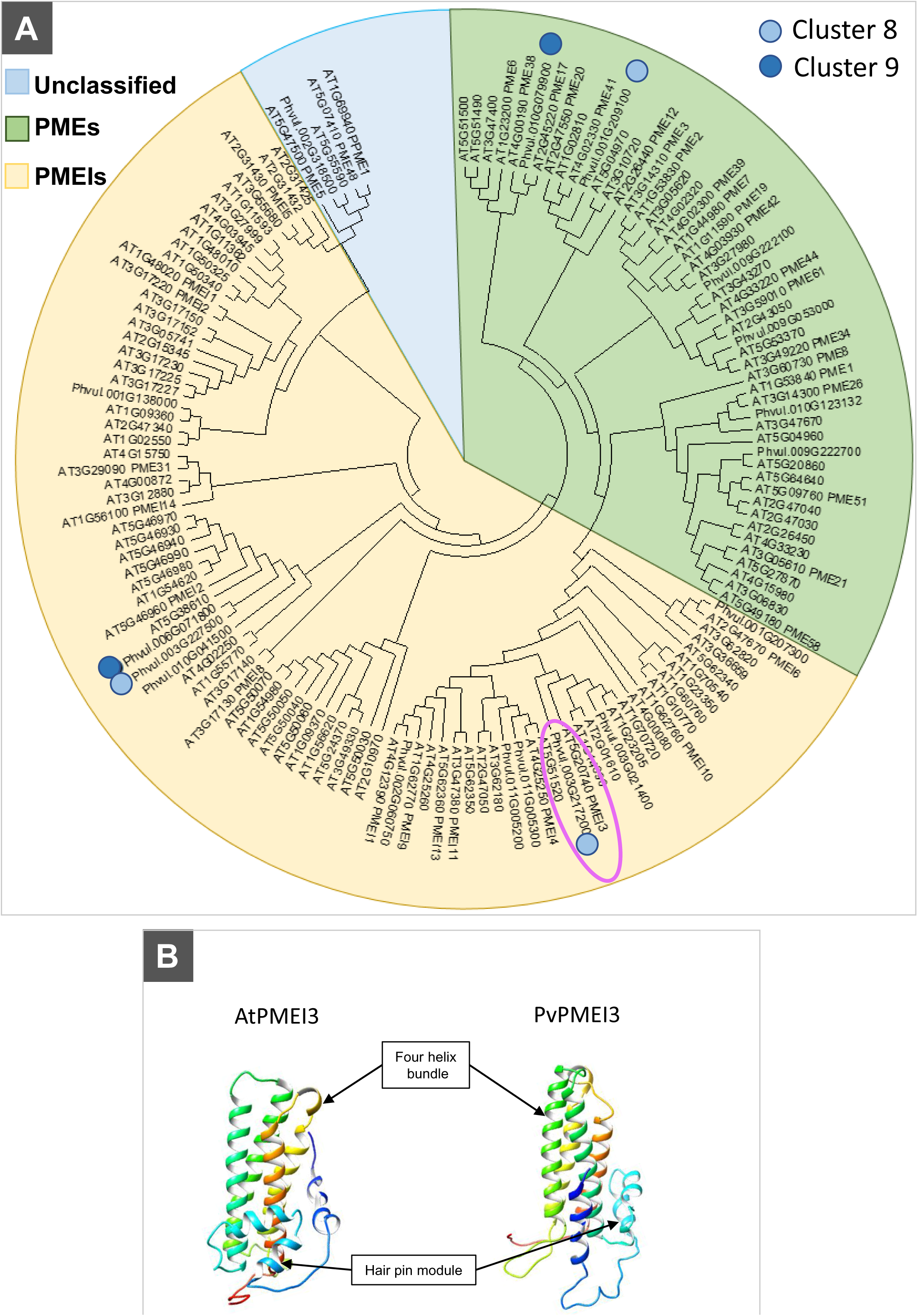
*Pseudomonas syringae* pv *phaseolicola* (Pph) produces an overexpression of several pectin methylesterase (PME) and pectin methylesterase inhibitor (PMEI) genes in common bean leaves. A. Phylogenetic analysis of common bean (Phvul.) and arabidopsis (AT) PME and PMEI proteins. The phylogenetic tree was obtained using the Arabidopsis PME and PMEI protein sequences, the protein sequences of all PME and PMEI genes deregulated by Pph infection, and those present in the RNA-seq clusterization. PMEs and PMEIs from cluster 8 and 9 were denoted with light blue and dark blue points, respectively. A common bean orthologue of AtPMEI3, the novel PvPMEI3 (Phvul. 003G217200), was selected to study its putative role in Pph infection (surrounded in pink). B. AtPMEI3 and PvPMEI3 3D protein models were obtained from primary sequences using the Chimera and Phyre2 program.

### AtPMEI3 plays a role in resistance against Pph

At the moment, there is limited information about the possible role of Arabidopsis PMEs and PMEIs in plant-pathogen interactions (Sénéchal et al., 2014; Bethke et al., 2014; Coculo and Lionetti, 2022). However, there seems to be a relationship between the modulation of HG methylesterification and Pph infection in common bean. To further evaluate this hypothesis, and considering the limitations of working with common bean, we tested Pph susceptibility in several *pme* and *pmei* mutants from Arabidopsis, including mutants in At*PMEI3.* All mutants tested affect PMEs and PMEI that were expressed in leaves (Supplementary Table 3; Winter et al., 2007; Yang et al., 2008). Leaves of these mutants were syringe infiltrated with Pph, using Arabidopsis Col-0 plants as negative controls because Pph is not able to infect this genotype (Xin and He, 2013).

Interestingly, 6 days after Pph infection, the appearance of a visible chlorosis in leaves was observed only in *pmei3-1*, *pmei13-1* and *pmei3-1xpmei13-1* double mutant (Figure 11A). Moreover, another independent experiment with two allelic mutants for both genes showed similar results (Supplementary Figure 8). The percentage of damaged areas was higher in *pmei3-1* mutant and *pmei3-1xpmei13-1* double mutants than in the single *pmei13-1* mutants (Figure 11B and Supplementary Figure 7). In the same way, the quantification of colony forming units (CFUs) from Pph-infected leaves was higher in *pmei3-1* and *pmei3xpmei13* (Figure 11C). These results suggest that AtPMEI3 and, to a minor extent, AtPMEI13 should play a role in the tolerance of Arabidopsis to Pph.

**Figure 11:**
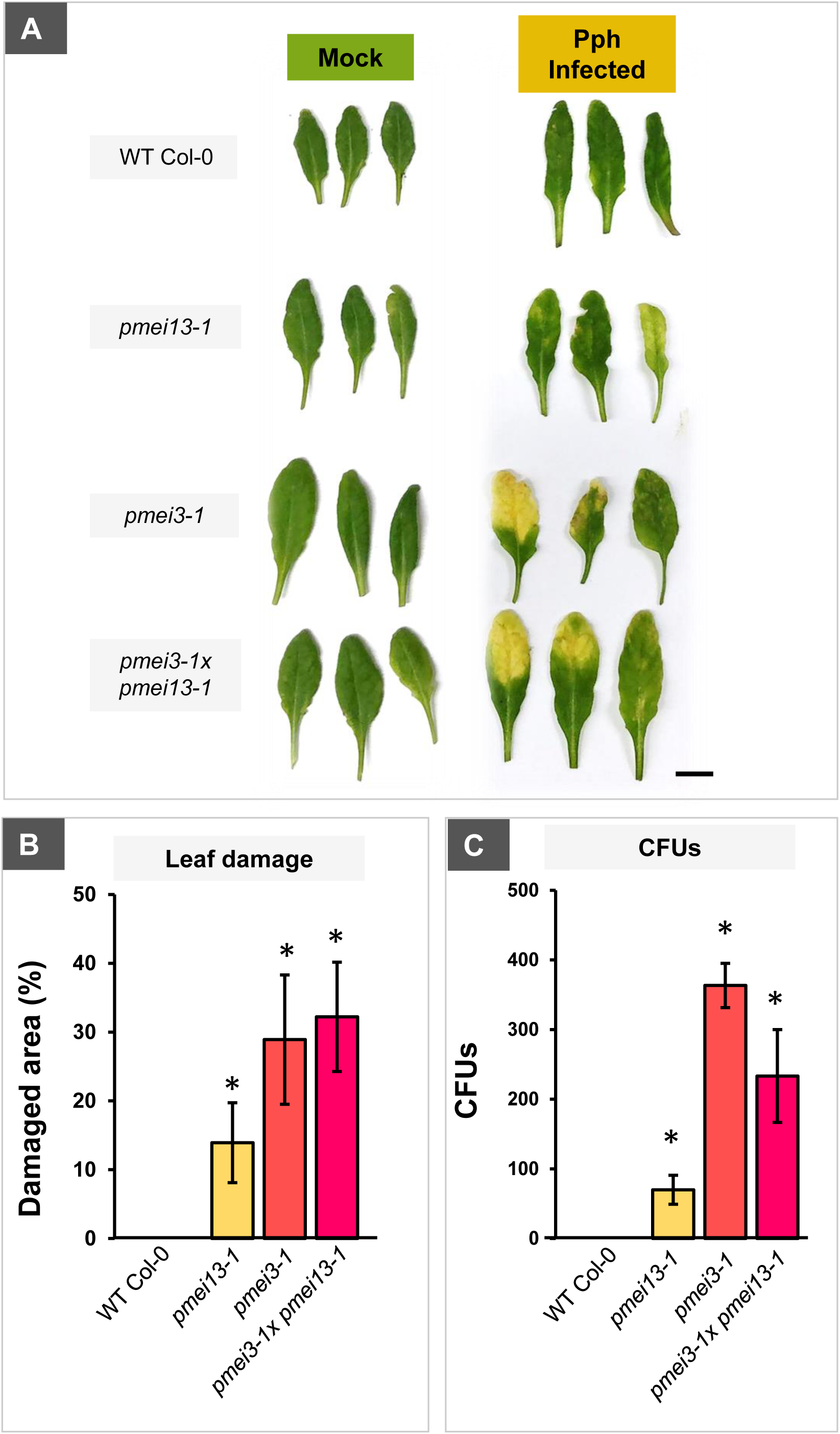
*Arabidopsis thaliana pmei3* and *pmei13* mutants showed susceptibility to *Pseudomonas syringae* pv *phaseolicola* (Pph) infection. A. Pph disease symptoms in *pmei13-1*, *pmei3-1* and *pmei3-1xpmei13-1* double mutants. Arabidopsis WT Col-0 and pmei mutants were grown for 6 weeks, and then, Pph was syringe-infiltrated in leaves. After 72 hours, infected and non-infected leaves were photographed. The infection caused visible damage in *pmei* mutants, especially in *pmei3-1*. Bar: 1 cm. B. Pph disease symptoms were quantified from WT Col-0 and *pmei* mutants leaves using imageJ v1.53c. Error bars show the SE from three biological replicates. Asterisks show significant differences using Student’s t-test (p<0.05). C. B. Pph Colony Forming Units (CFUs) in damaged leaves. Error bars show the SE from three biological replicates. Asterisks show significant differences using Student’s t-test (p<0.05).

### PvPMEI3 is a pectinmethylesterase inhibitor which is located in the apoplast

*PvPMEI*3 showed the highest expression in common bean leaves upon Pph infection (Supplementary Figure 6) and is the orthologue to At*PMEI13*, which could have a role in Pph infection (Figure 11). As a putative PMEI, this protein is expected to be located in the apoplast and have inhibitory activity against PMEs. To investigate this, a deeper characterization of PvPMEI3 was carried out (Figure 12).

**Figure 12:**
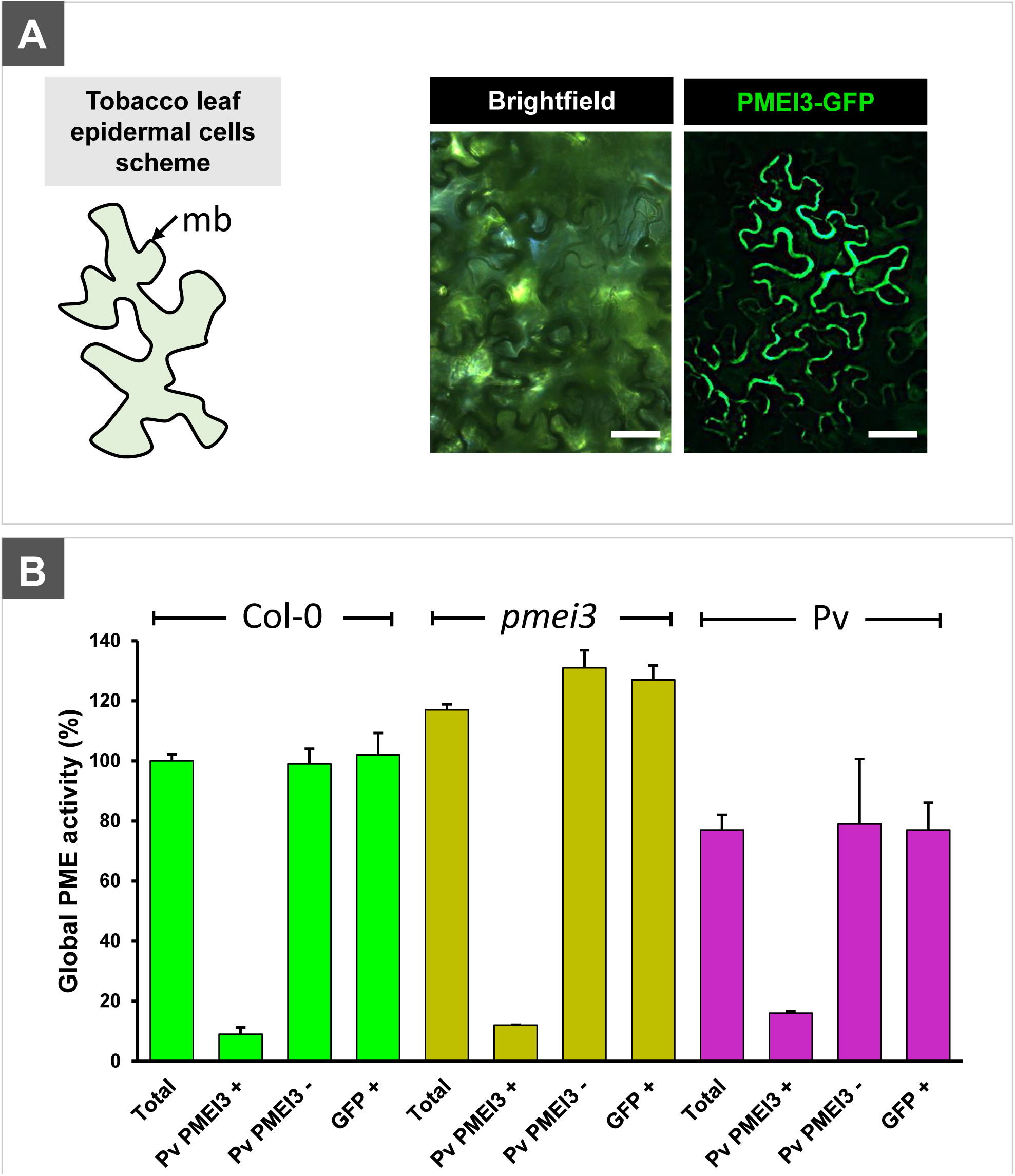
PvPMEI3 is located in the apoplast and is able to inhibit PME activity *in vitro*. A. Left panel: schematic representation of a typical *N. benthamiana* (tobacco) epidermal leaf cell. mb: cell membrane. Right panel: Subcellular localization revealed that PvPMEI3 protein is accumulated in the apoplast of tobacco leaves transformed with the CaMV35S-PvPMEI3-GFP construct. Bars: 25 mm. *B. In vitro* PME inhibitory activity of PvPMEI3. Escherichia coli was transformed with a vector carrying the T7-PvPMEI3-Hys(6x) construct. IPTG-induced (+) and non-induced (-) E. coli protein extracts were incubated with total protein extracts from Col-0, pmei3-1 mutant and common bean to evaluate the PME inhibitory activity of PvPMEI3. PME activity was calculated in comparison with the activity of Col-0 (100 %). Proteins from *E. coli* expressing 35S-GFP were used as negative control.

First, GFP fluorescence of overexpressed PvPMEI3-GFP protein in tobacco leaves showed that, as expected, this protein is mainly accumulated in the apoplast (Figure 12A), indicating that PvPMEI3 is a secreted protein that performs its action in the CW.

Second, PvPMEI3-Hys(6x) was cloned in *E. coli* under the control of an IPTG-inducible promoter. The total protein content from IPTG-induced and non-induced *E. coli* was extracted and used to determine its inhibitory activity. against common bean, Arabidopsis WT Col-0 and *pmei3-1* mutant proteins (Figure 12B). Proteins from *E. coli* expressing 35S-GFP were used as negative control. Interestingly, *pmei3-1* mutant plants showed an increment of approximately 15 % of PME activity compared to control (Figure 12B). The incubation with the PvPMEI3 extracted showed an almost complete depletion of PME activity in WT Col-0, *pmei3-1* and *P. vulgaris* protein extracts (Figure 12B). Moreover, non IPTG-induced and GFP (alone) *E. coli* protein extracts did not show changes in PME activity. These results suggest that PvPMEI3 can inhibit PME activity in Arabidopsis and common bean leaf protein extracts, being the first PMEI activity characterized in common bean.

## DISCUSSION

### Transcriptional reprogramming upon Pph infection would explain common bean susceptibility

In nature, plants must overcome multiple stresses, such as those induced by pathogens, by activating several defense mechanisms. However, many important crops are susceptible to pathogens, such as the common bean variety *riñón* used in this study, which is susceptible to Pph (Monteagudo et al., 2006; De la Rubia et al., 2021). Although several advances have been made to reveal the mechanisms behind this lack of resistance (De la Rubia et al., 2021a; De la Rubia et al., 2021b), the study of the changes in gene expression upon Pph infection of common bean leaves would help to understand the infection process. Therefore, we performed a transcriptomic analysis by RNA-seq of infected common bean leaves at 0, 2 and 9 hours post infection, once we collected evidence that the infection was taking place (Figure 1). Transcriptomic data revealed a misregulation of 1,575 DEGs when comparing Pph-infected and non-infected plants at 2 hours and, interestingly, most of the DEGs were found upregulated (1,076) (Figure 3.A). However, at 9 hours post infection the number of DEGs decreased to 565, indicating that the large reprogramming of gene expression due to Pph infection that takes place at 2 hours was being stopped at 9 hours. In comparison with other previous plant-pathogen transcriptomic studies, the number of genes deregulated were similar to those obtained in Arabidopsis against *P. syringae DC3000* (Truman et al., 2006), and in common bean after *Xanthomonas phaseoli* inoculation (Foucher et al., 2020). However, although there are cases where major misregulation of DEGs takes place when a susceptible common bean is attacked by *X. phaseoli* (Foucher et al., 2020), in the specific case of inoculation with an hypervirulent *P. syringae DC3000* strain (with *hrp* virulence gene), it induced less transcriptomic reprogramming than wild-type bacteria (Truman et al., 2006), suggesting a blockade at some points of the first defense lines. Some microorganisms, such as some *P. syringae* strains, are able to introduce effectors into the plant cell cytoplasm (Dillon et al., 2019), which disrupt signaling or defensive reprogramming of the plant (Sun et al., 2020). This stage, known as Effector Triggered Suppression (ETS), causes the development of the disease (Jones and Dangl, 2006). Cluster analysis of RNA-seq point to this type of ETS response, due to the fact that most of the DEGs were grouped in cluster 8 (326) and cluster 9 (890) and showed up-regulation at 2 hours of infection (suggesting PTI activation) followed by down-regulation at 9 hours post infection (when ETS is supposed to take place) (Figure 3.B). GO enrichment analysis of these two clusters showed a great number of genes related to immune and defense responses and their regulation (Supplementary Figures 2 and 3). Interestingly, this tendency was observed for a huge number of plant receptors that belong to families previously associated with plant immune responses (Supplementary Table 2). Therefore, these data supported the idea that Pph infection did not provoke a sustained induction of plant extracellular receptors and genes related to defense responses to avoid bacteria recognition, which, in the end, allowed Pph to infect common bean leaves.

A similar trend occurred with respect to SA-and JA-related genes (Supplementary Figures 2, 3 and 4), which was also confirmed by phytohormone quantification, as Pph infection provoked the highest accumulation of SA and JA at 2 hours, which was not sustained at 9 hours (Figure 4). As in other physiological processes, PHs play an important role during the immune defense activation and maintenance, although the events following pathogen recognition remain largely unknown (Couto and Zipfel, 2016). Among others, SA, JA, ABA and ethylene are required for different immune responses against different pathogens (Bari and Jones, 2009). SA has been shown to be a key player in the resistance against biotrophic pathogens (Zhang and Li, 2019), regulating the expression of defense-related genes, depending on its low, intermediate or high concentration, producing different responses such as hypersensitive response (HR) or systemic acquired resistance (SAR) (Zhang and Li, 2019). For different pathosystems, the negative cross-talk between JA and SA is well-known (Cristina et al., 2010) although, in Arabidopsis upon *P. syringae* infection, SA and JA levels increase once the pathogen effectors are recognized (Liu et al., 2016). In the same way, both PHs increased only at 2 hours post infection in our pathosystem, but probably not with the intensity and duration required for resistance. On the other hand, many pathogens inhibit PH production by means of effectors in order to be more successful in their pathogenesis (Katsir et al., 2008) and this could be the reason for the decrease of SA and JA at 9 hours post infection.

Finally, in both Cluster 8 and 9 some GO biological functions such as “response to oxidative stress” appear (Figure 3.B, Supplementary Figures 2 and 3), which could be associated with the reactive oxygen species (ROS) burst described upon pathogen attack (Couto and Zipfel, 2016). In *riñón* variety, the H_2_O_2_ production takes places at short times post infection but not with the extent and duration necessary for stopping the pathogen invasion (De la Rubia et al., 2021a), which is also in concordance with the expression profile of these oxidative stress genes. In addition, among others, the terms “response to water deprivation” or “stomatal closure” can be found in both clusters, which could indicate the transpiration reduction and water loss by closing stomata, which renders in restricting pathogen entry through stomatal pores (Bharath et al., 2021). Although no difference in the content of ABA was observed among infected and control plants in this experiment, the ABA-content profile of infected plants showed and attenuation with respect to control when the content of this PH is evaluated 24 hours after infection (De la Rubia et al., 2021a), indicating that Pph could be producing effectors for blocking the production of ABA and thus, avoiding stomatal closure.

### Pph infection provoked the quick, but not sustained, overexpression of common bean CW-related genes, accompanied by changes in homogalacturonan structure

The CW is the first plant defensive layer against intruders (Bacete et al., 2018). Plants are able to rearrange this structure on demand at different developmental stages. The CW can be modified by changing the proportions of its main components, and/or the type and extent of cross-linking among CW polymers (Bacete et al., 2018; Vaahtera et al., 2019). In this study, a great number of CW-related gene deregulations was detected, and many of them were grouped again in cluster 8 and 9 (Table 1). Several callose-related genes were found, all of them belonging to cluster 9 (overexpressed only at 2 hours post infection), being callose a β-glucan previously detected after *P. syringae* infections (Wang et al., 2021). Interestingly, the Pph infection also provoked a deregulation of CW-biosynthesis (glycosyltransferases and phenylpropanoid pathway) and CW-remodeling (Glycosyl hydrolases, PMEs and PMEIs) genes. This did not render significant changes at 2 hours post infection in monosaccharide quantification of the AIR upon infection (Figure 5.A). However, at 9 hours, an increase in Ara and a decrease in Xyl were detected (although differences in Rha and GalA were not found) suggesting changes in hemicelluloses (xylan, xyloglucan or arabinoxylan) and/or pectin side chains (RG-I and RG-II). Moreover, it seems that Pph did not affect total pectin amounts or, at least, HG and RG-I backbones, because the GalA (and Rha) did not show any significant change at any of the times evaluated (Figure 5.A). These results are in accordance with previous results obtained in common bean leaves with Pph infections at longer times, where the presence of Pph provoked changes in pectin extractability and other properties, such as the HG methylesterification degree (De la Rubia et al., 2021b). Interestingly, this increment in GalA content was accompanied by a higher methanol release from AIR at 9 hours after infection (Figure 5.B), which would suggest a higher methylesterification degree of HG. Immunolabelling of Pph-infected and non-infected common bean leaf sections with JIM7, JIM5 and 2F4 antibodies (which bind to highly and poorly methylesterified HG and egg-box structures, respectively), showed a higher labelling of all antibodies at 2 and 9 hours post infection, although the increment of labelling for JIM5 and 2F4 was confined in the parenchymatic tissues in the leaves. These areas with a higher level of JIM5 and 2F4 labelling could be associated with the points where Pph was proliferating (Figure 6 and 7). These results confirm that the Pph infection would not provoke a substantial and sustained deposition of newly synthesized pectic polysaccharides, but it changed pectin properties such as the degree of HG methylesterification, which seemed to take place in a tissue-dependent manner. This idea is supported by the overexpression of several genes related to pectic metabolism (PME and PMEI), which are grouped in clusters 8 and 9 (only overexpressed at 2 hours post infection, Table 1), suggesting the specific control of the degree, and possibly pattern, of pectin methylesterification during Pph infection. PME activity and the concomitant changes in pectin methylesterification have been previously related to immune processes of Arabidopsis against *Pseudomonas syringae* pv *maculicola* (Bethke et al., 2014). Our results could be indicating that Pph produces pectin modifying enzymes during the first steps of the attack to reach the plasma membrane and introduce its effectors into the cell in order to block the plant defense. In line with this, common bean plants primed with the SA analog INA show a modification in the pectin domains, which reduced the susceptibility to Pph (De la Rubia et al., 2021b). These results are also similar to those observed in Arabidopsis lines with reduced pectin content, *ERF014* RNAi and IDL6 overexpressed lines, which became susceptible to *P. syringae* DC3000 (Zhang et al., 2016; Wang et al., 2017).

### The importance of homogalacturonan metabolism-related genes in the Pph infection

As stated above, the Pph infection caused the overexpression, only at 2 hours post infection, of several pectin metabolism-related genes, which are mainly associated with the control of the HG methylesterification degree, such as PMEs and PMEIs (Table 1). It has been proposed that a higher HG methylesterification degree would protect the HG from being degraded by pectin metabolic enzymes, such as PGs and/or PLs from hemibiotrophic and necrotrophic pathogens, especially necrotrophic fungi (Lionetti et al., 2012 and 2015; Wormit and Usadel, 2018). Being Pph an hemibiotrophic pathogen, we wonder if this bacterium could secrete pectinases to degrade pectins from plant CWs. Pph reference genome encodes 1 PL and 2 PGs, according to database https://img.jgi.doe.gov/. In addition, the overexpression of one PL gene (*pnlA*) and one PG gene (PSPPH_A0072) have also been described when Pph was incubated with tissue extracts from a susceptible variety of *P. vulgaris* (Hernández-Morales, et al., 2009). PL and PG activities were present in the pelleted bacteria and secreted to the KB media (Supplementary Figure 5). Interestingly, when Pph was incubated in the presence of PGA, the activity of both proteins was higher in the culture media, indicating an increase in the secretion of these enzymes in the presence of unmethylated pectins. However, Pph did not show PME or PMEI activities, and neither putative PME/PMEIs were found *in silico* (Supplementary Figure 5). Taking all of this together suggests that Pph could secrete PL and PG enzymes into the apoplast to degrade pectins, facilitating its progress into the cytoplasm during the colonization process. Although these PG and PLs are more abundant in necrotrophic microorganisms such as fungi (Lionetti, 2015; Lionetti et al., 2017), which usually cause a massive CW degradation, bacteria also produce CW-degrading enzymes (CWDEs). However, the production of these pectin degrading enzymes could be a strategy to develop a fast colonization that allows the pathogen to introduce the effectors, using the type III secretion system, which block the plant immune system (Lo et al., 2017).

The presence of certain PMEs and/or PMEIs seemed to be essential to plant resistance to pathogens (Lionetti et al., 2007; 2012; Bethke et al., 2014; Wormit and Usadel, 2018; Corpo et al., 2020). For instance, the expressions of Arabidopsis *PME17* and *PMEI12* are necessary for the resistance against the necrotrophic fungus *Botrytis cinerea* (Lionetti et al., 2017; Corpo et al., 2020). In concordance to this, we observed the overexpression, at 2 hours post infection, of two common bean PME genes (Phvul-010G079900 and Phvul.001G209100), closely related to Arabidopsis *PME17*; and two PMEIs (Phvul.006G071800 and Phvul.003G041500), closely related to Arabidopsis *PMEI12* (Table 1; Figure 10.A). However, this expression profile was not maintained in time, which could also be one of the reasons why common bean is susceptible to Pph. In addition, we observed the overexpression of one PMEI (Phvul.003G217200) ortholog to PMEI3 from Arabidopsis (Table 1; figure 9.A). Considering these changes in pectinase gene expression, we decided to evaluate the activity of pectinases during the Pph infection. Interestingly, the analysis of PME, PG and PL activities showed a decreasing trend during the infection, which could protect the pectin structure at short times. However, this fact could also reduce the release of oligogalacturonides and methanol (from the demethylesterification of HG) during the infection, molecules that can be sensed by the plant to activate PTI responses (Ferrari et al., 2013). The decreasing amount of these molecules could also cause an increase in the susceptibility to Pph at longer times.

### Arabidopsis mutants affecting PMEIs are susceptible to Pph infection

PME activity is regulated, among other factors, by PMEI proteins, whose role in plant pathogenesis has been previously described (Bethke et al., 2014; Coculo and Lionetti et al., 2022). For instance, the infection of Arabidopsis with *P. syringae* pv. *macuicola* produces an enhancement of total PME activity, however the lack of specific PMEs causes a reduction in total PME activity and leads to higher susceptibility to infection with this pathovar (Bethke et al., 2014). Recently, Arabidopsis *pmei13* mutant has been described to have increased susceptibility against *Myzus persicae* aphid (Silva-Sanzana et al., 2019), indicating the potential role of AtPMEI13 in defense processes. In this work, we observed susceptibility to Pph infection in several *pme/pmei* Arabidopsis mutants (Supplementary Table 3). From all the mutants selected, only *pmei3* (Arabidopsis ortholog of *PvPMEI3*), *pmei3xpmei13* double mutant and, to a minor extent, *pmei13* showed visible damage at 6 days post infection with Pph (Figure 11.A and B; Supplementary Figure 7). In addition, *pmei3* mutants showed an enhancement in the PME activity in comparison with Col-0 (Figure 12.B), and both *pmei3* and *pmei13* have been previously described to have *in vitro* PMEI inhibitory activity and also to affect pectin methylesterification in Arabidopsis (Peaucelle et al., 2008; Silva-Sanzana et al., 2019; Xu et al., 2022). Taking all these results together, it seems that the specific modulation of HG methylesterification, together with changes in the distribution of HG domains with a difference in methylesterification, are key factors in resistance or susceptibility to Pph, at least, in common bean and Arabidopsis. It is also interesting that, besides the expression of *PvPMEI3*, Pph causes the overexpression of other possible *PMEI* and *PME* genes in common bean. These results open the door to deeper study of the role of PMEs and PMEIs in Pph infection. It also arises the question whether these overexpressed *PMEI*s could inhibit the overexpressed *PME*s or, in fact, inhibit other PMEs that could also play a role in defense, but whose expressions were not affected by the presence of Pph.

### PvPMEI3 is a novel apoplastic PME inhibitor with a possible role in the defense responses of common bean to Pph

As AtPMEI3 expression contributes to the resistance of Arabidopsis to Pph, we wondered if PvPMEI3 also has PMEI activity. As in other PMEIs (Coculo and Lionetti, 2022), the GFP fluorescence of PvPMEI3-GFP protein in tobacco leaves revealed an apoplastic accumulation of this protein (Figure 12.A).

Moreover, the presence of PvPMEI3-GFP resulted in a reduction of PME activity in all leaf extracts tested, indicating that PvPMEI3 is actually a PMEI with a possible role in defense, similar to other pectin-modifying enzymes that have been described to participate in this process (Lionetti et al., 2017; Silva-Sanzana et al., 2019; Corpo et al., 2020; Xu et al., 2022). The discovery of this novel common bean PMEI activity and the important role of the modulation of HG methylesterification open the possibility to explore new biotechnological approaches focusing on CWs to develop defense strategies for common bean against Pph.

In this study, we analyzed the transcriptome and the CW modifications at 2 and 9 hours after Pph infection in 15-day-old common bean plants. RNA-seq analysis showed an overexpression of defense-related genes at 2 hours, but the expression profile was not sustained throughout the infection. This trend was also observed in the quantification of defense-related PHs, such as SA, JA and ABA, and in CW-related gene expression. The CW monosaccharide analysis did not reveal differences with respect to control at 2 hours post infection, and changes in arabinose and xylose at 9 hours, suggesting that the Pph infection produced modifications in matrix polysaccharides. Apart from this, the main pectin modifying activities (PL, PG and PME) decreased upon infection, while the expression of genes which encode inhibitory activities, such as PG inhibitors and PMEIs, was increased. As a result, the HG methylesterification degree increased, and immunolabelling assays showed an increment of HG with high and low methylesterification degree, together with the formation of more egg-boxes, indicating the important role of HG in the infection process. These results are very interesting and considering that Pph showed *in vitro* PG and PL activities, the exploration of the secretion and action of these bacterial enzymes should be addressed in the future. In addition, Arabidopsis *pmei3* and *pmei13* mutants showed a susceptibility to Pph that control plants did not show, reinforcing the idea that the modulation of HG methylesterification is a key factor in resistance against Pph. Finally, in this work, it was demonstrated that PvPMEI3 is an apoplastic protein with PMEI activity, making this gene a good novel biotechnological target to develop future defense strategies for common bean against Pph.

## METHODS

### Plant materials and growth conditions

Common bean (*P. vulgaris* L.) seeds cultivar *riñón*, were obtained from the Protected Geographical Identification in La Bañeza-León region (Spain). Once sterilized with 70% (v/v) ethanol for 30 s and 0.4% (w/v) NaClO for 20 min, the seeds were washed with sterile water and placed to germinate *in vitro* on universal sterilized substrate (Blumenerde, Gramoflor, made in Germany). Plants were grown in growth chambers at 25 ± 2°C with a 16 h photoperiod under a photon flux density of 45 ± 5 μmol m^−2^ s^−1^ provided by daylight fluorescent tubes (TLD 36W/830, Philips), as indicated in De la Rubia et al. (2021b).

Arabidopsis (*Arabidopsis thaliana*) wild-type Col-0, *pme and pmei* mutants (Supplementary Table 3) were obtained from ABRC (http://abrc.osu.edu/) using the SIGnAL Salk collection (Alonso et al., 2003). WS4 and *pmei13-2* (FLAG_58B07) were kindly donated by Dr. Helen North (INRA Versailles). Plants were grown in growth chambers at 22 ± 4 °C with a 16 h photoperiod under a photon flux density of 100 ± 5 μmol m^−2^ s^−1^, 65% relative humidity, using TopCrop^®^ compost as a substrate, and watered every week with TopCrop^®^ liquid growth fertilizer, as indicated in Saez-Aguayo et al., (2021). Tobacco (*Nicotiana benthamiana*) plants were grown under the same conditions.

### Bacteria growth conditions and plant inoculation

*Pseudomonas syringae* van Hall 1902, the *P. syringae* pv. *phaseolicola* 1448A from CECT321 (Pph) was grown at 30 °C on liquid King’s B (KB) medium with shaking at 220 rpm until an optical density 600 (OD_600_) of 0.8 was obtained.

For infection experiments, a final Pph concentration of 10^8^ CFU/mL was used. The Pph solution for common bean inoculation was prepared removing the media by centrifugation at 6000 rpm for 10 minutes and the pellet was resuspended in sterile water with 0.1% (v/v) Tween 20 as indicated in De la Rubia et al., (2021a). Inoculation was achieved by spraying at least 2 mL of the Pph solution on the upper and lower side of cotyledonary leaves of common bean plants at developmental stage V1 (approximately 14 days post-germination, when the two cotyledonary leaves were completely expanded). For Arabidopsis *pmei* mutants, infection was developed at 30 days post-germination by infiltrating, at least, 0.5 mL Pph solution in sterile 10 mM MgCl_2_ *per* leaf.

### Tissue collection, RNA isolation and RNA sequencing

As observed in the experimental design figure (Figure 1A), the cotyledonary foliage leaves of 10 plants *per* treatment were collected at 0, 2 and 9 hours post-infection, and they were homogenized with liquid nitrogen to make a pool for each condition. All homogenates were stored at −80°C until their use and three independent experiments (n = 3) were performed. The RNA was isolated using RNeasy Plant Micro kit (Qiagen) and TURBO^TM^ DNAse (Invitrogen), following the manufacturer’s instructions. The quality and concentration of total RNA samples were analyzed in a Nanodrop and had at least a 260/280 nm ratio of 1.8. In addition, RNA quality numbers were obtained using an Agilent 2100 Expert Bio-analyzer (from Inbiotec Institute, Universidad de León). Samples were sent to “Centro Nacional de Análisis Genómico” CNAG where, from 1.0 μg of total RNA, cDNA libraries for each sample were sequenced using the TruSeq^®^ Stranded mRNA-seq kit (Illumina) by next generation sequencing (NGS), under the repository PHASEOLUS_01.

### DAB staining and PCR for checking Pph infection in bean leaves

To check the damage produced by the Pph infection in leaves, the 3,3′-diaminobenzidine (DAB) staining protocol of Daudi and O’Brien (2016) was followed with little modifications. Infected and uninfected leaves were incubated 4 hours in fresh DAB staining solution (1mg mL^-1^ DAB, 10 mM Na_2_HPO_4_) in dark and shaking conditions. Then, the DAB solution was replaced by bleaching solution (ethanol: acetic acid: glycerol; 3:1:1; v/v/v) and samples were placed in boiling water for 15 minutes to eliminate all chlorophyll. The area of damaged tissue was measured using Fiji software (Schindelin et al., 2012).

To detect the presence of Pph in infected samples, a PCR was performed using the same homogenized tissue as in the RNAseq experiment, using the specific primers PphFW (5’-3’: GACGTCCCGCGAATAGCAATAATC) and PphRev (5’-3’: CAACGCCGGCGCAATGTCG) employed by Seok Cho et al. (2010), at 10 μM final concentration. The amplification cycle consisted of 15 min at 96°C, 40 cycles of 15 seconds at 52°C and 30 seconds at 72°C, ending with a final elongation phase of 1 min at 72°C, in a T-Gradient Thermal Cycler (Applied Biosystems). Amplifications were visualized in 1.5 % agarose electrophoresis developed at 80V for 75 min.

### RNA sequencing analysis

Raw data reads (.fastq files) were trimmed using the Trim Galore Cutadapt (Martin *et al*., 2011) to remove adapters and N’s, filtering by quality (minimum Q30) and length (minimum 50 bp). Trimmed reads were aligned against the *Phaseolus vulgaris* V2.1 reference genome (DOE-JGI and USDA-NIFA, assembly of the *Phaseolus vulgaris* G19833 genome) using STAR (Dobin et al., 2013) with default parameters. SAMTools were used to sort by name (sort.bam using the function-n) (Li et al., 2009). Mapped read pair counts were calculated with FeatureCounts function (Liao et al., 2014) in R (R Development Core Team, 2020). After filtering all sequences that did not map at least with one gene in all variables (low abundance filtering with FeatureCounts function in R), an analysis of differentially expressed genes (DEGs) was performed using DESeq2 (Love et al., 2014), considering a false discovery rate (FDR) of <0.05. Up-regulated and down-regulated genes were used to develop a co-expression analysis with coseq function (Rau and Maugis-Rabusseau, 2018) in R, using Model *K*means. To identify enriched Gene Ontology terms, the enrichGO function was used (Yu et al., 2016) in R, using FDR and Qvalue < 0.05, pAdjustMethod BH, and *A. thaliana* TAIR10 as background (Arabidopsis Genome Initiative, 2000). Finally, results were visualized using ggplots2 and ‘heatmap.2’ function of R statistical package.

### Phytohormone determination by LC/MS

PHs were extracted following the protocol described by de la Torre-González et al., (2017) with some modifications (De la Rubia et al., 2021). 150 mg FW of homogenized tissue were used for these analyses. The chromatographic conditions and the MS/MS parameters were those optimized and described in Supplementary Table 4. The MassLynxTM software (version 4.1, Waters) was used. The quantification was conducted by means of standard curves (1, 2.5, 5, 10, 25, 50, 75, 100, 200 and 500 ng/mL of ABA, JA and SA, Sigma Aldrich).

### Alcohol-insoluble residue (AIR) extraction

A volume of homogenized tissues equal to that used for RNA-seq analysis was washed in 10 volumes of the following organic solvents, as indicated in Saez-Aguayo et al., (2021). Firstly, three washes of 80 % (v/v) ethanol for 4 hours at room temperature (RT) were followed by washing the pellets twice with methanol/chloroform 1/1 (v/v) (3 h, RT). Then, two washes with acetone were carried out for 1 hour at RT to remove lipids. Finally, the residue was dried overnight, also at RT.

### Acid hydrolysis and monosaccharide determination by HPAEC-PAD

The AIR was hydrolyzed with 400 μL of 2 M trifluoroacetic acid (TFA) with 20 μL 5 mM callose and 20 μL 5 mM myo-inositol, as internal standards, for 1 hour at 121°C, as indicated in Parra-Rojas et al., (2019) with little modifications. Samples were dried under nitrogen stream at 40°C and washed twice with 100% 2-propanol, to be dried under nitrogen again afterwards. The products were resuspended in 1 mL of water, filtered (pore size: 0.45 mm) and used for HPAEC-PAD analysis immediately.

The hydrolyzed products were analyzed in a Dionex ICS3000 ion chromatography system, which was equipped with a pulsed amperometric detector, a CarboPac PA1 (4×250 mm) analytical column and a CarboPac PA1 (4×50 mm) guard column, following the Parra-Rojas et al., (2019) method.

### Methylesterification analysis

For measuring methanol in AIR samples, 1mg AIR was saponified with 1M NaOH and the released methanol was measured as described by Saez-Aguayo et al., (2013). All experiments were developed using five technical replicates and the three biological replicates that conformed the samples from the RNA-seq analyses.

### Determination of PME, PL and PG activities

The pectin methyesterase (PME) activity was determined as indicated in Sáez-Aguayo et al., (2013), with little modifications. Relative PME activity was determined using 50 μg proteins in 20 μL which were loaded into 6 mm diameter wells, in gels prepared with 0.1 % (w/v) of pectin from apple (with 50-70 % methyl-esterification, Sigma-Aldrich), 1 % (w/v) agarose, 12.5 mM citric acid, and 50 mM Na_2_HPO_4_ pH 6.5. Plates were incubated at 28°C overnight and PME activity was determined as described in Saez-Aguayo et al., 2013 using Fiji software (Schindelin et al., 2012) and considering Col-0 as 100 % of activity. The pectate lyase (PL) activity was determined following the indications of Marín-Rodríguez et al. (2003). The protein extraction was performed using 10 mM Tris-HCl pH7, 1 mM EDTA and 100 mM NaCl extraction buffer. The reaction mixture was prepared separating solution A (120 mM Tris-HCl pH 8.5, 0.48 % polygalacturonic acid; w/v) and solution B (0.6 mM CaCl_2_) to prevent the Ca^2+^ used as cofactor by PLs from forming egg-boxes with polygalacturonic chains. Reactions were started with 150 μL solution A, 150 μL solution B and 30 μL protein extraction (200 μg protein). Then, absorbances were measured at 235 nm after 1 hour of stabilization (Time 0) and after 15 hours at 30°C (Time 15). Results were expressed as the change of absorbance between T15 and T0, indicating the release of products with double bonds from polygalacturonic acid as a substrate.

To determine the polygalacturonase (PG) activity, proteins were extracted with the PME extraction buffer described above. In this case, 200 μg of total proteins was assayed following the indications in Silva-Sanzana et al. (2019). For this, 0.1 mL protein extracts were assayed with 0.3 mL of reaction buffer (200 mM NaCl and 200 mM sodium acetate, pH 4.5) and 0.3 mL of 1% (w/v) polygalacturonic acid for 60 minutes at 37°C. The reactions were stopped by heating at 100°C for 5 minutes. Then, 100 μL of the reaction were mixed with 100 μL DNS reagent and heated at 100°C for 30 minutes. Samples were cooled on ice, 1 mL of milli-Q H_2_O was added and absorbances at 540 nm were measured. A standard curve of Galacturonic acid (0.2-10 μg/ml) was used to quantify the PG activity.

For determining PME, PMEI, PL and PG activity in Pph, the bacteria were grown in different media: KB, KB + 0.5 % PGA (Sigma; w/v), KB + 0.5 % apple pectin (Sigma; w/v) or KB + 0.5 % apple pectin + 1 U Pectinesterase (Sigma), testing the supernatant and the pellet (both extracted with extraction buffer), and Pph growth.

### Immunolabeling

The infection of common bean leaves was performed as described above and distal parts of uninfected and infected leaves were fixed following the method described by Balic et al. (2014). Tissues were incubated with a series of increasing Neo-Clear^®^ (Merck) concetrations (from 30 % to 100 %; v/v in 100 % ethanol). Tissues were immediately embedded in ParaPlast^®^ (Sigma-Aldrich) in two steps: firstly in 50 % ParaPlast^®^/Neo-Clear^®^ (v/v) and secondly in 100 % ParaPlast^®^. Three-μm microtome sections were obtained to perform immunolabeling with three monoclonal antibodies: JIM5 (poorly methylesterified HG), JIM7 (highly methylesterified HG) and 2F4 (egg-box structures). After removing the ParaPlast^®^, the sections were blocked by incubation in 2 % milk in TBS (w/v) for 30 min at RT. Afterwards, they were incubated with primary antibodies (JIM5 and JIM7 at 1:50 and 2F4 1:100 in 2 % milk in TBS-Tween 20) for 2 hours. After incubation, the samples were washed three times with TBS-Tween 20 followed by incubation with 1:1000 secondary antibody in 2 % milk in TBS-Tween 20 for 1.5 hours (Anti-rat IgG for JIM5 and JIM7 and Anti-mouse IgG for 2F4 couples to Alexa-Fluor; Molecular Probes). The sections were washed three times with TBS-Tween 20 and then incubated with propidium iodide (20 mg ml^−1^) for 10 min. Finally, they were mounted in 90 % glycerol in TBS (v/v) to obtain optical sections using a Leica TCS LSI spectral confocal laser scanning microscope as indicated in Parra-Rojas et al., (2019).

### Phylogenetic analysis of PME/PMEIs and structural prediction of PvPMEI3

Five putative *P. vulgaris* PME/PMEIs overexpressed upon Pph infection were selected to construct a phylogenetic tree, together with all the PME/PMEIs found in Arabidopsis TAIR database (https://www.arabidopsis.org/). All the predicted protein sequences encoded by these genes were aligned using Muscle v3.8, (Edgar, 2004). After that, a maximum likelihood phylogeny, following the Dayhoff matrix based model, was inferred with MEGA7 (Kumar et al., 2016).

Regarding structural prediction, the protein sequences were analyzed with **P**rotein **H**omology/analog**Y R**ecognition **E**ngine V 2.0 Phyre^2^ (http://www.sbg.bio.ic.ac.uk/phyre2/html/page.cgi?id=index) and the putative structure was visualized with UCSF Chimera (Pettersen et al., 2004).

### Cloning procedures

Total RNA was extracted from common bean (*P. vulgaris*) leaves using RNeasy Plus Mini Kit (Qiagen). After DNase I (ThermoFisher Scientific) treatment, RNA was used as a template to synthesize cDNA using the retro-transcriptase Superscript II (Invitrogen) following the manufacturer’s directions. PvPMEI3 full-cds was cloned with and without the stop codon by using the primers PMEI3Fw (5’-3’: ATGGCAACAAAGCTACTGTGGC) and PMEI3StopRv (5’-3’: TCACGGCGTCTGAGTAGTGC) PMEI3NoStopRv (5’-3’: CGGCGTCTGAGTAGTGCGGTATT). The products were purified using E.Z.N.A.^®^ Gel extraction kit (Omega Bio-Tek) and inserted into a pCR8 vector, according to standard protocols (Life Technologies), obtaining the pCR8-PMEI3-Stop and pCR8-PMEI3-NoStop vectors. After sequence verification of all clones, the PMIE3-NoStop clone was fused at C-terminal with green fluorescence protein (GFP) by recombining the pCR8-PMEI3-NoStop vector with the pGWB505 vector, using LR clonase (ThermoFisher Scientific). Finally, vector pGWB505-PMEI3-NoStop was obtained. This construction allowed the expression of PMEI3-GFP under the control of 35S promoter.

Moreover, for PMEI activity determination, PMEI3 was fused to His(6x) at N-terminal by the recombination of pCR8-PMEI3-Stop vector with pET300 vector, to obtain the pET300-PMEI3-Stop vector. This construction allowed the expression of His(6x)-PMEI3 under the control of an *Escherichia coli* IPTG-inducible promoter.

### *In vivo* subcellular PvPMEI3 localization

To analyze the PvPMEI3 protein localization, pGWB505-PMEI3-NoStop vector was transformed into A*grobacterium tumefaciens* strain GV3101 (Holsters et al., 1978). After growing *A. tumefaciens* until OD_600_ 0.4 in LB media, bacteria were pelleted by centrifugation and resuspended in sterile 10 mM MgCl_2_ + 20 μM acetosyringone. After two hours of incubation at RT, 30-day-old tobacco leaves were infiltrated with *A. tumefaciens,* carrying the vector pGWB505-PMEI3-NoStop, with a syringe. Finally, images from the lower epidermis of transformed leaves were taken after 48 hours of infection with an epifluorescence microscope Olympus IX2-UCB coupled to a MIcroPublisher 3.3 RTV camera. The argon laser excitation line was set to 488 nm to detect the GFP fluorescence. Tobacco leaves infiltrated with a GFP construct without PMEI were used as appropriate control.

### PvPMEI3 activity determination

The pET300-PMEI3-Stop vector was used to transform *Escherichia coli* Rosetta (DE3), which was growth overnight in LB media with 100 μg/mL ampyciline at 220 rpm and 37°C. Then, PMEI3 expression was induced by adding 0.1 mM IPTG for 5 hours at 220 rpm 37°C. After that, the induction media was removed by centrifugation and proteins were extracted by the resuspension of bacteria in B-PER™ Bacterial Protein Extraction Reagent (ThermoFisher Scientific) with 0.8 U/mL DNase I (ThermoFisher Scientific), 40 μg/mL Lysozyme, 40 μM PMSF, following the manufacturer’s instructions. *A. tumefaciens* protein extracts from the supernatant were used to analyze the PMEI activity.

For PMEI activity analysis, proteins from Col-0 and *pmei3-1* mutant line were extracted with the PME extraction buffer, and protein quantification was determined as described above. The analysis of PMEI activity was measured in PME activity agarose plates, by the inhibition of the halo appearance by the mixture of Col-0, *pmei3-1* and common bean protein extracts with the PvPMEI3 protein extract.

### Expression analysis by quantitative real-time PCR (qRT-PCR)

Expression of selected genes were analyzed by qRT-PCR to validate the RNA-seq analysis. The analysis was developed as described in De la Rubia et al. (2021a). The housekeeping gene used in this study was *Ukn1* (Borges *et al*. 2012) and all data were analyzed using the 2^-ΔΔCt^ method (Livak, and Schmittgen, 2001). The primers of the genes selected for RNA-seq validation (Supplementary Table 4), except for PR1 (Guerrero-González et al., 2011), were designed using Primer-BLAST (Ye et al., 2012).

The data from DEseq normalized counts were statistically compared with those from qRT-PCR by Pearson’s correlation in GraphPad Prism 8 (GraphPad Software, San Diego, California USA, www.graphpad.com) using the default parameters.

### Statistical analysis and data representation

All results were statistically analyzed by GraphPad Prism 8 (GraphPad Software, San Diego, California USA, www.graphpad.com). The respective statistical test, as well as the number of replicates, is indicated in each caption. Significant statistical differences were indicated in each graph with an asterisk (*p*<0.05).

## Supplemental Figure Legends

**Supplementary Figure 1: Heatmap of the top-100 most differentially expressed genes (DEGs) for three biological replicates (R1, R2, R3) of the RNA-seq analysis.** The treatments were control at 0 hours (C_0), control (C_2) and infected plants at 2 hours (I_2), and control (C_9) and infected plants at 9 hours (I_9).

**Supplementary Figure 2: Enrichment of gene ontology (GO) terms using DEGs of Cluster 8.** A high number of genes are related to immune responses to pathogens. Defense-related GO terms are highlighted in yellow boxes and, in green, those associated with phytohormone responses.

**Supplementary Figure 3: Gene ontology (GO) terms enrichment of Cluster 9 DEGs.** A high number of genes are related to immune responses to external biotic stimuli. Defense-related GO terms are highlighted in yellow boxes and, in green, those associated with phytohormones.

**Supplementary Figure 4: DEGs listed in the GO term** “response to salicylic acid” (GO:0009751) (A) and “response to jasmonic acid” (GO:0009753) (B) of cluster 8.

**Supplementary Figure 5: Pectin-related enzymatic activities in *Pseudomonas syringae* pv. *phaseolicola* (Pph).**

A. Polygalacturonase activity in Pph grown in KB media with glycerol (KB) or with polygalacturonic acid (PGA) as carbon source. The activity was measured in the supernatant (media) and the pelleted bacteria (bacteria) for both media. Values show media and ES from three biological replicates (n=9).

B. Pectin lyase activity in Pph grown in KB media with glycerol (KB) or with polygalacturonic acid (PGA) as carbon source. The activity was measured in the supernatant (media) and the pelleted bacteria (bacteria) for both media. Values show media and ES from three biological replicates (n=9).

C. PME inhibitory activity of all Pph extracts explained above, which were incubated with comercial PME (Sigma). The results were expressed compared to a comercial pectinesterase (100 %). Values show media and ES from three biological replicates (n=9).

**Supplementary Figure 6: Gene expression patterns from the RNA-seq analysis for the putative PvPMEIs:** A. Phvul.003G217200.1 (PvPMEI3), B. Phvul.003G227500.1 and C. Phvul.006G071800.1. The average expression profiles are the normal counts obtained after differential expression analysis using DESeq2. Error bars represent SE values from three biological replicates (n=9).

**Supplementary Figure 7: AtPMEI3 and PvPMEI3 protein alignment.** The alignment was developed using CLUSTAL O (1.2.4) multiple sequence alignment. The two sequences have an identity of 38.07 % and a query coverage of 83 %. The colors show the different aminoacid classes: In RED, Small (small+ hydrophobic (incl.aromatic-Y)); in BLUE, Acidic; in MAGENTA, Basic – H; and in GREEN, Hydroxyl + sulfhydryl + amine + G.

An asterisk (*) indicates positions which have a single, fully conserved residue. A colon (:) indicates conservation between groups of strongly similar properties. A period (.) indicates conservation between groups of weakly similar properties.

**Supplementary Figure 8: *Arabidopsis thaliana pmei3* and *pmei13* mutants showed susceptibility against *Pseudomonas syringae* pv. *phaseolicola* (Pph) infection.**

A. Pph disease symptoms evaluated in *pmei13*, *pmei3* and p*mei3-1xpmei13-1* double mutants at 6 days post infection. Bar: 1 cm.

B. Leaf damage area quantification caused by Pph infection. The appearance of the typical yellow damage patterns associated with Pph infection was quantified using ImageJ program.

## Supplemental Tables

**Supplementary table 1. Quality metrics of libraries** for comparative transcriptomic experiments for the three biological replicates (R1, R2, R3) of control plants at time 0 (Control 0h), and control and infected plants at 2 hours (Control 2h and Infected 2h) and 9 hours (Control 9h and Infected 9h) post infection.

**Supplementary Table 2. Differentially expressed plant receptors** related to immune responses

**Supplementary Table 3. Analysis of leaf-expressed *pme* and *pmei* Arabidopsis mutant plants with a putative role in *Pseudomonas syringae* pv. *phaseolicola* (Pph) infection.**

PMEs and PMEIs were selected considering their absolute expression in vegetative rosettes and mesophyll leaves and guard cells of 5-week-old (5WO) plants obtained from eFP browser (Winter, et al., 2007; Yang et al., 2008). Highest absolute gene expression values were shaded in blue for comparison. The absolute expressions of defense-related *PME17* and *PME41* genes were also included for comparison.

**Supplementary Table 4. Optimized mass spectrometry parameters** for the quantitation of target phytohormones and quantitation results under optimized UPLC-MS/MS conditions.

**Supplementary Table 5. Sequence of the primers** used to validate the RNA-seq analysis by qRT-PCR.

## Author Contributions and Acknowledgments

P.G-A., A.L-G., S.S-A. and A.G. designed the research. A.G., A.L-G., R.Y., P.S-O, D.S. performed the experiments. ML.C., A.G. performed the phytohormone analysis, C.M and A.R analyze transcriptomic data. P.G-A., A.L-G., S.S-A., A.G and C. M. analyzed the data. A.G., A.L-G., P.G-A. and S.S-A., wrote the article. All authors have read and agreed to the published version of the manuscript. We thank Rafael Calvo for the assistance with the English manuscript correction.

